# UbiB proteins mediate an ATP-dependent decarboxylation step in bacterial ubiquinone biosynthesis

**DOI:** 10.64898/2026.03.05.709815

**Authors:** Katayoun Kazemzadeh, Bruno Faivre, Sophie-Carole Chobert, Sophie Saphia Abby, Julie Michaud, Tuan Anh Dinh, Chilin Alexandre, Lerouxel Olivier, Marc Fontecave, Pierrel Fabien, Murielle Lombard, Ludovic Pelosi

## Abstract

Polyisoprenoid quinones such as ubiquinone (UQ) play an essential role in cellular physiology, acting as membrane-bound electron and proton carriers in respiratory chains and other biological processes across all domains of life. In *Escherichia coli*, the canonical UQ biosynthesis pathway is well characterized. It involves twelve proteins (UbiA-UbiK and UbiX), most of which catalyzing one of the eight modifications of the aromatic ring derived from 4-hydroxybenzoic acid (4-HB), while others (UbiB, UbiJ, UbiK) act as accessory factors ensuring efficient UQ production. Following prenylation by UbiA and subsequent decarboxylation by the UbiX/UbiD system, the final six reactions are catalyzed within a soluble Ubi-complex. UbiB, an atypical protein kinase-like enzyme, was proposed to extract decarboxylated intermediates from the membrane and mediate their delivery to the Ubi-complex.

In this study, we demonstrate the existence of an alternative decarboxylation system in *E. coli*, as UQ biosynthesis can proceed in the absence of the UbiX/UbiD system. Our results show that this alternative decarboxylation activity depends on UbiB and requires its ATPase activity. Bioinformatic analyses further revealed that approximately 27% of *Pseudomonadota* species lack UbiX/UbiD homologs, and we found that UbiB proteins from two such species enhance the alternative decarboxylation activity when expressed in *E. coli*. In addition, we identified conserved residues in UbiB that are specifically required for decarboxylation but dispensable for the delivery of UQ intermediates to the Ubi-complex. Taken together, our findings support a model in which UbiB acts as an ATP-dependent decarboxylase, thereby broadening the functional scope of this poorly characterized protein.

**Importance:** Polyisoprenoid quinones, including ubiquinone (UQ, also known as coenzyme Q), are essential electron and proton carriers in respiratory chains across all domains of life. After prenylation and decarboxylation, the final six steps of UQ biosynthesis are carried out by a soluble Ubi-complex. The only decarboxylation system currently identified in the UQ pathway, the UbiX/UbiD system, is absent from numerous bacterial genomes, and no isofunctional enzymes have been described to date. UbiB, an atypical protein kinase-like enzyme, has been proposed to mediate the extraction of UQ precursors from the membrane, thereby rendering them accessible to the soluble Ubi-complex. Here, we show that UbiB proteins fulfill an additional function in bacteria. Specifically, they contribute to the decarboxylation of UQ precursors through a mechanism that remains unknown but depends on ATP hydrolysis. Given that UbiB is conserved in all UQ-producing bacteria, it may represent a widespread decarboxylation system within this essential metabolic pathway.

## Introduction

Isoprenoid quinones are central to bioenergetics, as they constitute obligate electron and proton carriers in the respiratory chains of organisms belonging to the three domains of life. The main isoprenoid quinones are naphthoquinones (menaquinone (MK) and demethylmenaquinone (DMK)) and ubiquinone (UQ), which is also called coenzyme Q in eukaryotes (1). (D)MK and UQ share the same polyisoprenoid tail but differ by the structure of the redox-active head group, namely a naphthalene ring and a benzene ring, respectively, and in the value of their redox midpoint potential (1). (D)MK are considered anaerobic quinones, since they function primarily in anaerobic respiration, whereas UQ is considered an aerobic quinone, because it supplies electrons mostly to the reductases that reduce O_2_ (2). The polyisoprenoid tail varies in length between organisms (UQ_6_ in *Saccharomyces cerevisiae*, UQ_8_ in *Escherichia coli*, and UQ_10_ in humans) and confers extremely hydrophobic properties to isoprenoid quinones, which are consequently localized in cellular membranes.

Over a period of a few decades, the UQ biosynthesis pathway was extensively studied in *E. coli*, and has emerged as a paradigm for the synthesis of UQ in oxic conditions and more recently in anoxic conditions (3). The O_2_-dependent UQ biosynthesis pathway involves twelve proteins (UbiA to UbiK and UbiX), while the O_2_-independent pathway involves ten proteins (UbiA to UbiE, UbiG, UbiU to UbiT and UbiX) (4). Most of these proteins catalyze modifications of the aromatic ring of the 4-hydroxybenzoic acid (4-HB) precursor and some of them are shared between the two pathways (reactions catalyzed by UbiA to UbiE, UbiG and UbiX). The first step in the biosynthesis of UQ is catalyzed by UbiC, which converts chorismate to 4-HB. Following its prenylation by UbiA, the phenyl ring of 4-HB is decarboxylated by the UbiX/UbiD system (Fig. 1A), which consists of the decarboxylase UbiD and its associated flavin prenyltransferase UbiX that produces the prenylated FMN indispensable to UbiD (5). The resulting decarboxylated intermediate is then further modified within a soluble complex (hereafter called Ubi-complex) (6) (Fig. 1A). Inside the Ubi-complex, the *S*-adenosyl-L-methionine-dependent UbiG and UbiE proteins catalyze *O*- and *C*-methylation reactions, respectively, while the O_2_-consuming flavoprotein monooxygenases (hereafter called FMO) UbiI, UbiH and UbiF, each hydroxylate one carbon atom of the aromatic ring (C5, C1 and C6, respectively) (7–10). The only reactions that differ between the O_2_-dependent and O_2_-independent pathways are the hydroxylation steps. Indeed, we have demonstrated that UbiU and UbiV catalyze these steps under anaerobic conditions using prephenate as the OH group donor, thus replacing Ubi-FMOs (4, 11, 12). In addition, UbiB, UbiJ, UbiK and UbiT exhibit supporting functions (3). UbiJ and UbiT share sequence identity and are both SCP2 (Sterol Carrier Protein 2) domain-containing proteins. The role of UbiT is limited to anoxic conditions (4, 12), whereas UbiJ, which interacts with UbiK (13–15), is important for UQ biosynthesis only in oxic conditions, playing a role in the assembly and/or the stability of the Ubi-complex (6). Finally, UbiB was originally assigned to the C5-hydroxylation step (16), which is now known to be catalyzed by UbiI (10). The UbiB family comprises bacterial UbiB and eukaryotic homologs Coq8 (ACDK3) and belongs to the unorthodox protein kinase-like family (17, 18). UbiB proteins from human, yeast and *E. coli* exhibit *in vitro* ATPase activity, which is stimulated by compounds that resemble UQ biosynthesis intermediates (18). Overall, as previously suggested in (18) and (3), UbiB family members are currently proposed to couple the hydrolysis of ATP to the extraction of decarboxylated UQ precursors from the membrane in order to make them available for UQ biosynthesis enzymes (Fig. 1A). However, this role as well as the molecular mechanism involved remain hypothetical.

**Figure 1:**
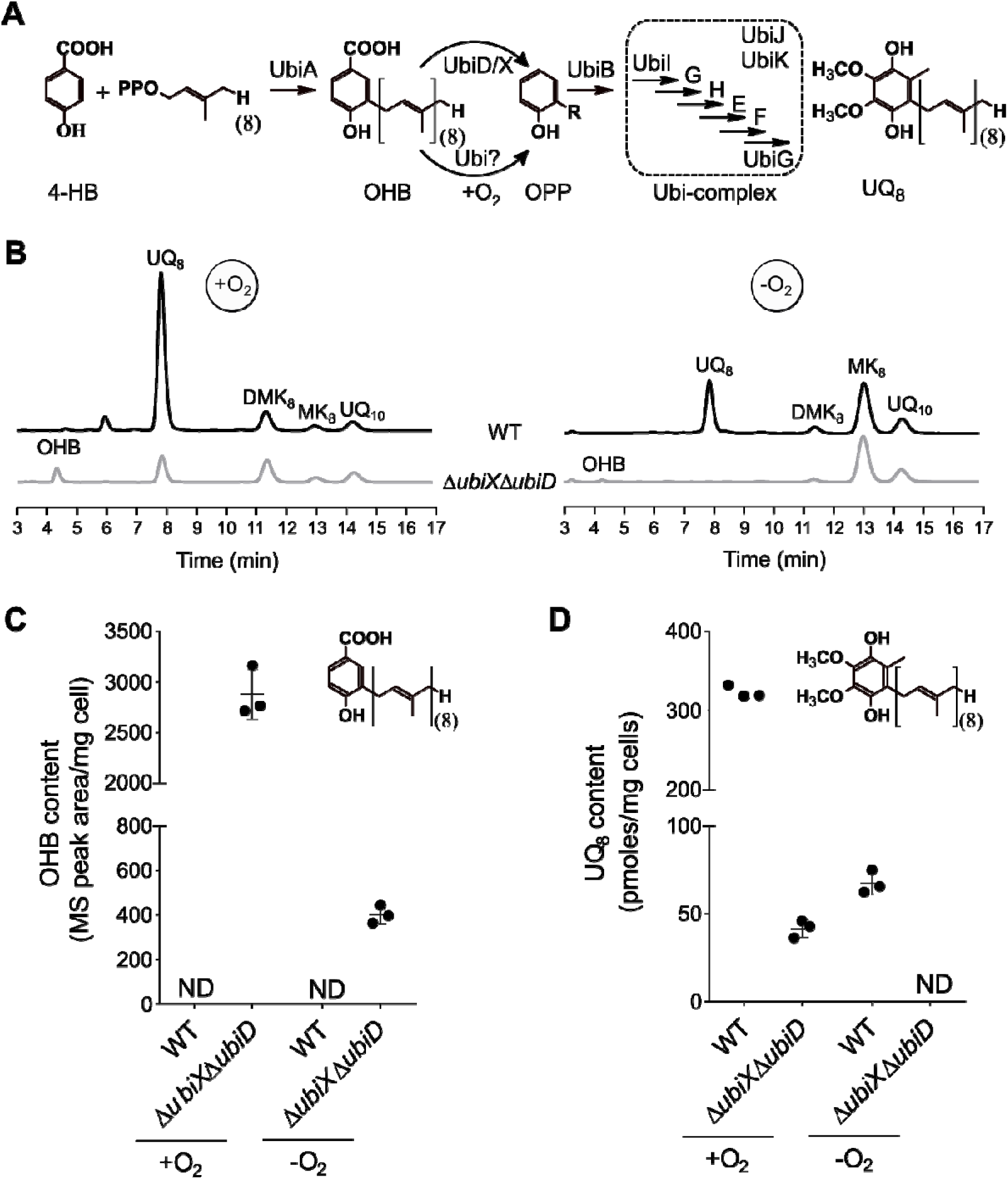
Aerobic UQ_8_ biosynthesis is maintained at low levels in the *E. coli* Δ*ubiX*Δ*ubiD* mutant strain. **(A)** Aerobic UQ_8_ biosynthesis pathway in *E. coli* and putative alternative decarboxylation step. OPP would be exported from the membrane to the soluble Ubi-complex by UbiB to be modified in UQ_8_. **(B)** HPLC-ECD analysis of lipid extracts from 1 mg of *E. coli* MG1655 (WT) and Δ*ubiX*Δ*ubiD* cells grown aerobically (+O_2_) or anaerobically (-O_2_) in LB medium. The chromatograms are representative of results from three independent experiments. The peaks corresponding to OHB, UQ_8_, DMK_8_, MK_8_ and UQ_10_, as a standard, are indicated. **(C-D)** OHB content (MS peak aera, mass detection M-H with m/z 681.3, C) and UQ_8_ content (ECD detection, D) in cells grown either aerobically (+O_2_) or anaerobically (-O_2_) in LB medium. The structure of each quantified compound is shown. Values are shown with means ± SD (n=3). Proteins belonging to the Ubi-complex are in framed. Abbreviations: OHB, octaprenyl-hydroxybenzoic acid, UQ_8_, ubiquinone 8, DMK_8_, demethylmenaquinone, MK_8_, methylmenaquinone and UQ_10_, ubiquinone 10. ND: not detected.

The current view of UQ biosynthesis in bacteria is mostly based on the *E. coli* pathway (3). However, studies combining computational and experimental approaches revealed an unsuspected diversity in the protein composition of the UQ biosynthesis pathway across bacteria, especially for the hydroxylation steps (19–21). The absence of the UbiX/UbiD system in UQ biosynthesis pathways that contain most other Ubi proteins has also been reported in various bacterial clades, suggesting that other enzyme systems may be involved in the decarboxylation of the prenylated 4-HB (22–25). In the present study, we discovered in *E. coli* a new decarboxylase involved in UQ biosynthesis, which is independent of the UbiX/UbiD system and functional exclusively under oxic conditions. Our results combining genetic, bioinformatic and biochemical approaches provide a strong link between UbiB and the new decarboxylase activity. We propose that UbiB, which is ubiquitous in UQ-producing bacteria (24), possesses the capacity to decarboxylate 4-hydroxy-3-polyprenylbenzoic acid into 2-polyprenylphenol in *Pseudomonadota* (formerly *Proteobacteria*).

## Results

### *E. coli* maintains some UQ_8_ biosynthesis in aerobic conditions in cells lacking the UbiX/UbiD system

In *E. coli*, the UbiX and UbiD proteins form a tightly linked system, which catalyzes the decarboxylation of octaprenyl-hydroxybenzoic acid (OHB), synthesized by UbiA, into octaprenylphenol (OPP) (26, 27). However, as already demonstrated by Gulmezian et *al*., inactivation of either the *ubiX* or *ubiD* genes leads to residual synthesis of UQ in *E. coli* (28). These results suggest the existence of an alternative OHB decarboxylation system in this bacterium. To confirm this, we analyzed an *E. coli* strain in which both the *ubiX* and *ubiD* genes were inactivated (Δ*ubiX*Δ*ubiD* mutant strain). Quinone content was analyzed by HPLC coupled to electrochemical detection (ECD) (Fig. 1B). Under aerobic conditions, the chromatogram of the Δ*ubiX*Δ*ubiD* mutant exhibited a peak at approximately 4.3 minutes (Fig. 1B, left panel). MS analysis of this peak revealed a predominant adduct (M-H) at m/z 681.3 corresponding to OHB (monoisotopic mass 682.5), consistent with a defect in the decarboxylation step (Fig. S1 and Fig. 1C). UQ_8_ was also detected at levels which amounted to 15-20% of those observed in wild-type (WT) cells (Fig. 1D), suggesting that OHB is still decarboxylated to some extent and subsequently converted into UQ_8_ through a mechanism independent of the UbiX/UbiD system. Interestingly, UQ_8_ was not detected in Δ*ubiX*Δ*ubiD* cells grown under anaerobic conditions (Fig. 1B, right panel and Fig. 1D), unlike OHB (Fig. 1C). Altogether, these results demonstrate that UbiX and UbiD are essential for *E. coli* to synthesize UQ_8_ under anaerobic conditions, whereas an alternative system enables the decarboxylation of OHB under aerobic conditions. We noted that inactivation of either the *ubiX* or *ubiD* genes resulted in the same phenotype as the double mutant (Fig. S2), as previously published (28).

### The *E. coli* alternative decarboxylase depends on O_2_

We investigated the role of O_2_ in the alternative decarboxylation step that compensates for the UbiX/UbiD system. We hypothesized that the alternative decarboxylase might be expressed exclusively under oxic conditions, might require O_2_ for activity, or both simultaneously. To explore this, we shifted a Δ*ubiDc* strain from anoxic to oxic conditions with or without prior chloramphenicol treatment to inhibit protein synthesis. The UQ_8_ content was monitored over a 14-hour period after the shift (Fig. 2A) (29). Prior to chloramphenicol treatment, UQ_8_ was not detected, and only OHB accumulated as previously shown in the Δ*ubiDc* strain (Fig. S2). UQ_8_ was detected shortly after the shift to oxic conditions, and its content gradually increased over 4 hours, regardless of chloramphenicol treatment. After 4 hours, the UQ_8_ level was approximately 60% of the level measured after overnight growth (Fig. 2B and Fig. S3). As expected, chloramphenicol inhibited cell growth of the Δ*ubiDc* cells throughout the experiment (Fig. 2C). Taken together, these results suggest that the alternative decarboxylase is expressed in *E. coli* under anaerobic conditions and that only the absence of O_2_ prevents its function.

**Figure 2:**
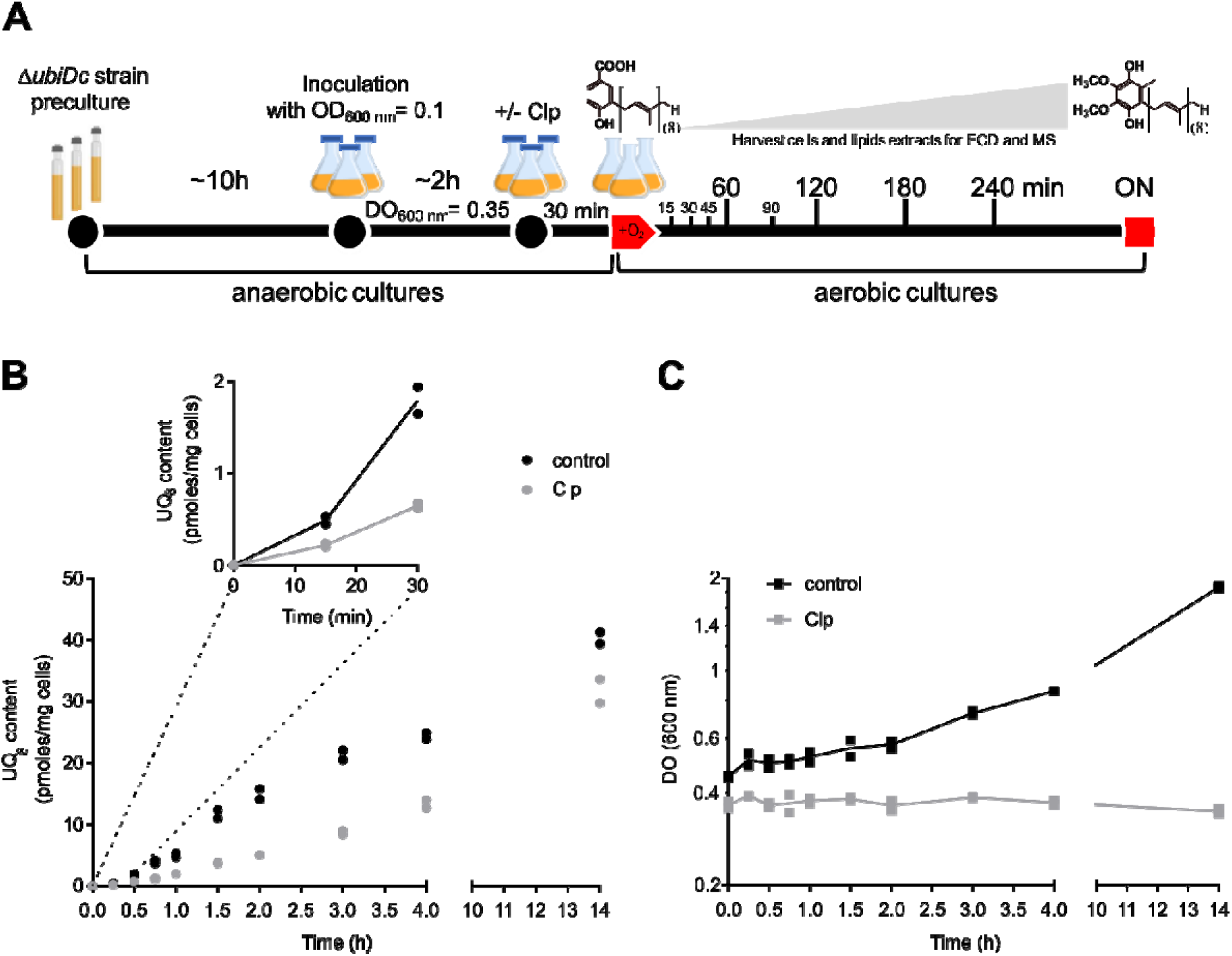
O_2_ is essential to the alternative decarboxylation of OHB. **(A)** Schematic representation of the experimental transition from anoxic to oxic conditions in the Δ*ubiDc* strain, showing the restoration of UQ_8_ biosynthesis from OHB, with or without (control) chloramphenicol (Clp) pretreatment. **(B)** Quantification of cellular UQ_8_ levels (MS peak aera, mass detection M+NH_4_^+^, m/z 744.5) after 4 hours and following overnight (ON∼14h) incubation. The inset highlights the UQ_8_ content restoration during the first 30 minutes of the experiment. **(C)** Growth curves of the Δ*ubiDc* strain cultured in LB medium under the same conditions than in (B). Data are representative of two independent experiments.

### UbiX/UbiD-independent UQ biosynthesis pathways are common in aerobic Pseudomonadota

To assess the distribution of UbiX and UbiD in UQ-producing bacteria, we leveraged our recent work on the quinone repertoire in *Pseudomonadota* (24) and queried publicly available genomes. Of the 3,928 species with genetic potential to produce UQ (presence of the core UbiABEG), 1,119 species (∼28%) lacked UbiX and UbiD homologs (Table S1). They were distributed as follows: 676 in *Alphaproteobacteria* (60.5%), 118 in *Betaproteobacteria* (10.5%), and 325 in *Gammaproteobacteria* (29%) (Table S1). These results indicate that decarboxylases alternative to the UbiX/UbiD system are likely widespread among UQ-producing *Pseudomonadota*. The presence of the O_2_-independent UQ biosynthesis pathway that we discovered a few years ago (4), is defined by the simultaneous presence of the genetic potential for the corresponding UbiU, UbiV and UbiT (UbiUVT) proteins (24). Whereas UbiUVT proteins were found in 39% of the total genomic dataset, they were identified only in 22 of 1,119 (∼2%) genomes that lacked UbiX and UbiD. In addition, none of them genomes harbored a complete (D)MK biosynthetic pathway, which is primarily associated with anaerobic metabolism (24). These findings suggest a link between aerobic metabolism and the decarboxylases alternative to the UbiX/UbiD system.

To investigate this, we used the metabolic information collected in our previous study (24), which covered ∼21% of the 3,928 species in our dataset, and significant associations were observed (Fig. 3). A high proportion (∼89%) of species lacking UbiX and UbiD, UbiUVT, and the (D)MK pathways was annotated as having an aerobic metabolism. The presence of UbiX and UbiD corresponded to an increased proportion of microaerophilic metabolism (∼9% versus ∼4%), and the additional presence of UbiUVT to an increased proportion of facultative metabolism (∼25% versus ∼7%). Finally, species with (D)MK pathways in addition to the UQ pathway showed a high proportion of facultative aerobic metabolism (∼51%). Collectively, these results suggest that most of the UQ-producing *Pseudomonadota* lacking both UbiD and UbiX rely primarily on aerobic respiration. This aligns with our biochemical findings, which highlight the essential role of O_2_ for the activity of the alternative decarboxylase.

**Figure 3:**
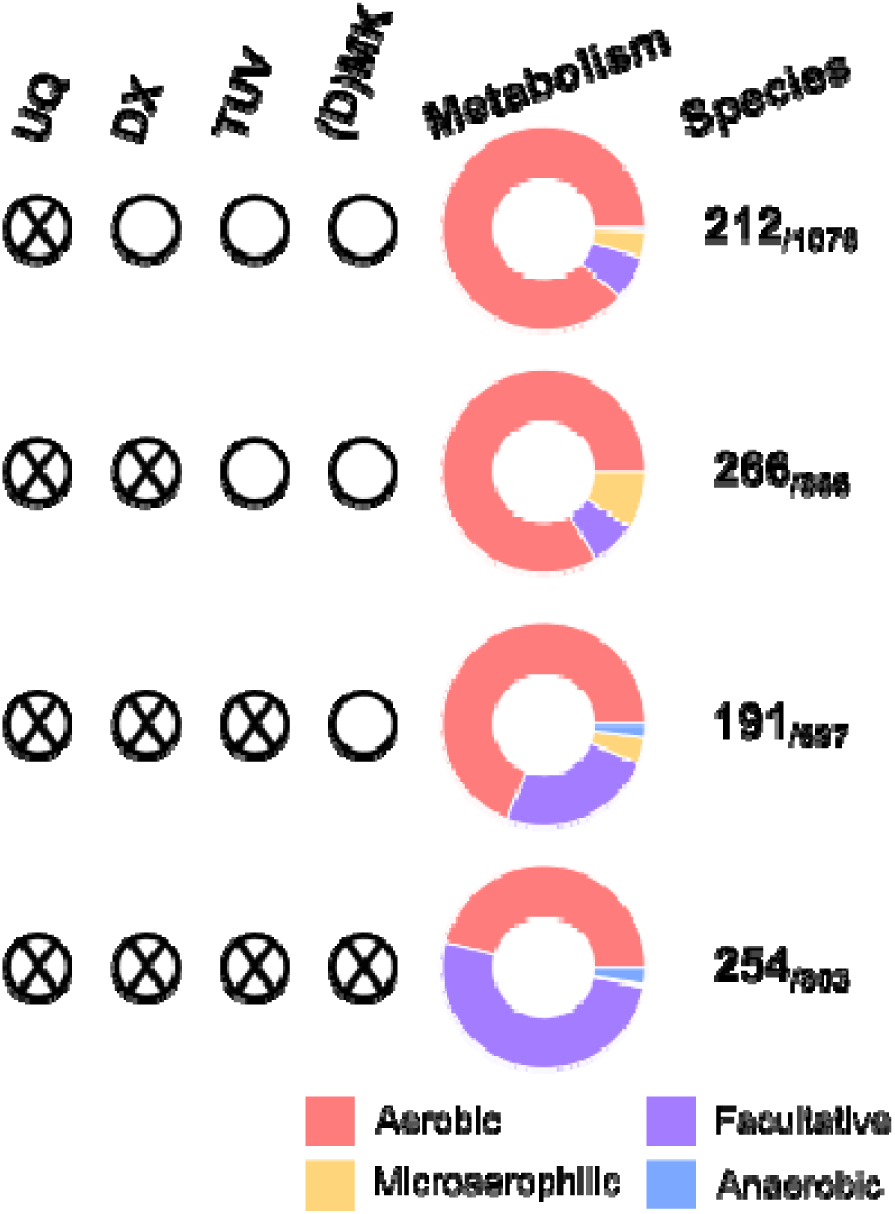
UQ biosynthesis pathways lacking the UbiX/UbiD system are widespread in aerobic *Pseudomonadota*. For each set of genes (DX, TUV) and pathways (UQ and (D)MK) (which presence is symbolized by a cross inside a circle, empty circle representing absence), the proportion of each metabolism (aerobic, facultative, microaerophilic, anaerobic) is shown. The column “Species” contains the fraction of species for which metabolic information was collected out of the total number of genomes screened.

Finally, an analysis of negative co-occurrence with UbiD and UbiX did not identify any alternative decarboxylase candidate. One possible explanation is that multiple enzymatic systems capable of decarboxylating 3-polyprenyl-4-hydroxybenzoic acid may coexist within a single bacterial species. Indeed, such functional redundancy was recently demonstrated in *Rhodobacter capsulatus*, in which the decarboxylation step can be carried out by the UbiX/UbiD system or by a novel class A flavoprotein monooxygenase, which has no homolog in *E. coli* (21).

### UbiB is required for the alternative OHB decarboxylation in *E. coli*

The UQ_8_ biosynthesis pathway in *E. coli* occurs in two stages during which UbiB could potentially play a pivotal role (3). Specifically, at the membrane level, UbiA prenylates 4-HB using octaprenyl diphosphate as a side chain precursor to generate OHB, which is predominantly decarboxylated by the UbiX/UbiD system to form OPP (Fig. 1A). Subsequently, UbiB is believed to extract OPP from the membrane, making it accessible to the soluble Ubi-complex for further modifications as proposed previously (3). Supporting this model, UbiB from *E. coli* copurified with early UQ biosynthesis intermediates like OPP and OHB and exhibited ATPase activity (18). To investigate whether the alternative decarboxylation of OHB occurs upstream or downstream of UbiB, we examined *E. coli* mutant strains in which the *ubiB* gene had been inactivated in wild-type or Δ*ubiX* genetic backgrounds (Δ*ubiB* and Δ*ubiX*Δ*ubiB*, respectively). As anticipated, the Δ*ubiB* mutant strain exhibited a strong decrease in UQ_8_ content compared to the WT strain, and accumulated large amounts of OPP without a trace of OHB, suggesting an efficient decarboxylation by the UbiX/UbiD system (Fig. 4A and Fig. 4B). In contrast, the Δ*ubiX*Δ*ubiB* mutant strain accumulated only OHB (Fig. 4C), while Δ*ubiXc* cells continued to synthesize UQ_8_, albeit at low level (Fig. 4A). Collectively, these findings are consistent with a decarboxylation step from OHB to OPP mainly carried out by the UbiX/UbiD system, independently of UbiB. However, when the UbiX/UbiD system is inactivated, UbiB becomes essential for the decarboxylation of OHB and, consequently, for UQ_8_ biosynthesis. Based on these results, we propose that the alternative decarboxylation of OHB to OPP is either directly facilitated by UbiB or occurs downstream of UbiB, i.e., at the level of the Ubi-complex.

**Figure 4:**
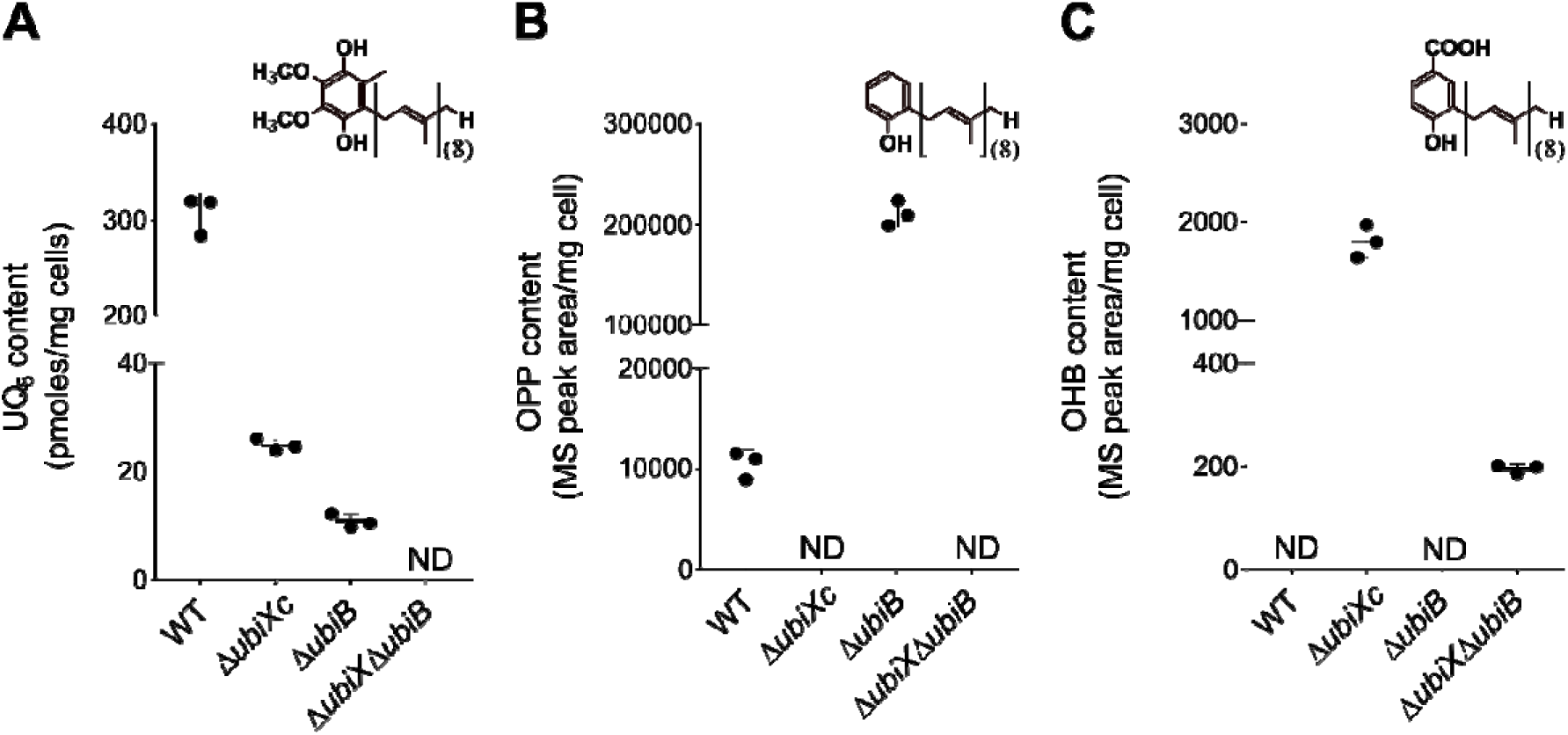
UbiB is essential to the decarboxylation of OHB when the UbiX/UbiD system is disabled. **(A)** UQ_8_ (ECD detection), **(B)** OPP (MS peak area, mass detection M+NH_4_^+^, m/z 656.5) and **(C)** OHB (MS peak aera, mass detection M-H, m/z 681.5) contents of lipid extracts from *E. coli* WT strain MG1655 and Δ*ubiXc*, Δ*ubiB* and Δ*ubiX*Δ*ubiB* mutant strains. Values are shown with means ± SD (n=3). The structure of each compound quantified is shown. ND: not detected.

To test this, we assessed the ability of UbiB and its D310N mutant (UbiB^D310N^), which exhibits a defect in ATPase activity (18), to restore UQ_8_ biosynthesis in the Δ*ubiB* and Δ*ubiX*Δ*ubiB* mutant strains. We observed that UbiB^D310N^ successfully restores UQ_8_ biosynthesis in the Δ*ubiB* mutant strain, reaching ∼70% of the UQ_8_ levels obtained upon expression of WT UbiB (Fig. 5A). In contrast, UbiB^D310N^ was unable to support UQ_8_ biosynthesis in the Δ*ubiX*Δ*ubiB* cells, whereas the expression of WT UbiB restored a UQ_8_ levels comparable to that of the Δ*ubiXc* cells (compare Fig. 4A and Fig. 5B). Therefore, we conclude that the D310 residue is dispensable for the canonical function of UbiB, as assessed in Δ*ubiB* cells, yet is strictly required for the alternative decarboxylation activity observed in Δ*ubiX*Δ*ubiB* cells.

**Figure 5:**
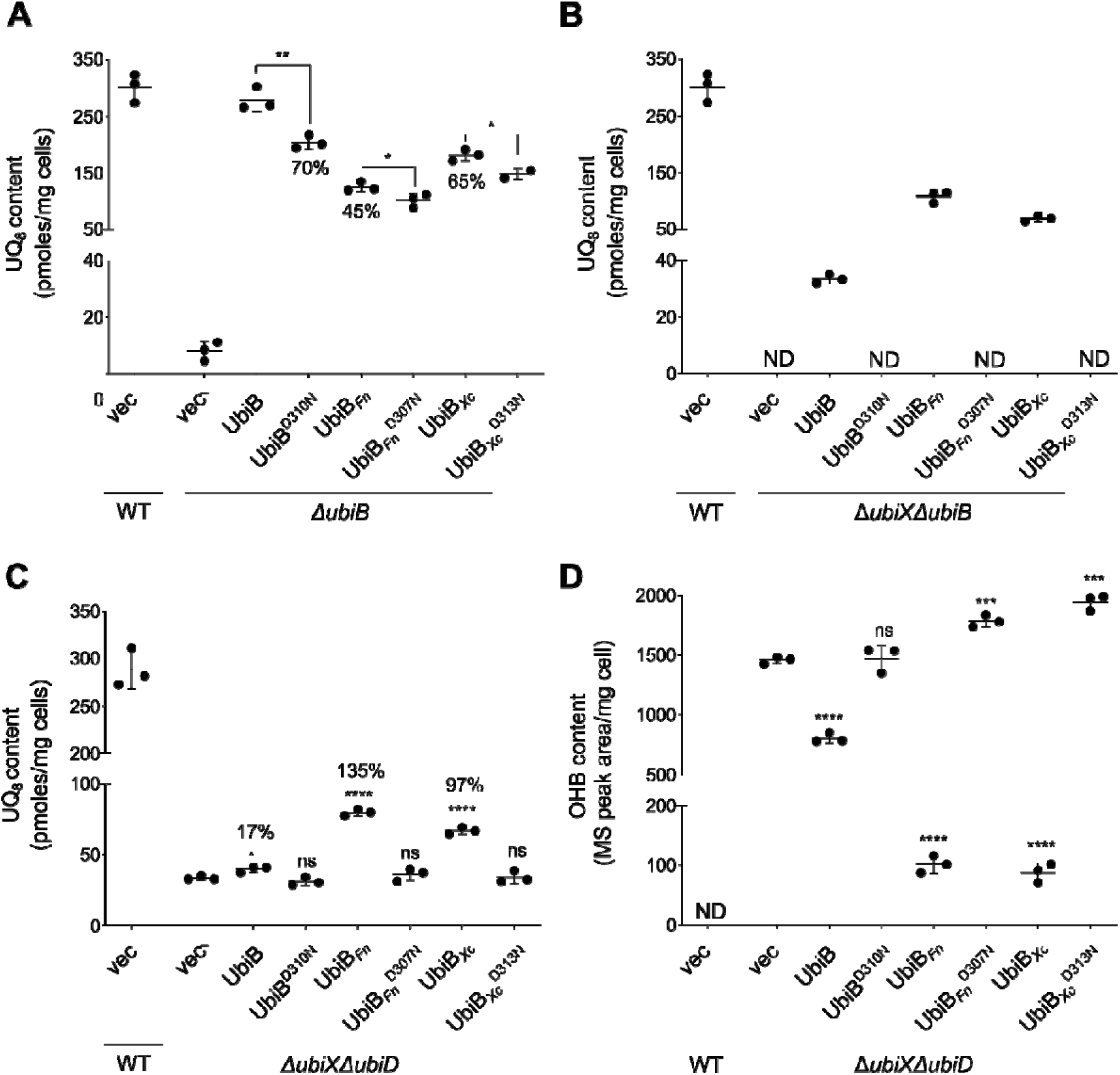
UbiB proteins exhibit an *in vivo* decarboxylase activity. WT UbiB, UbiB*_Fn_* from *F. novicida* and UbiB*_Xc_* from *X. campestris* and their respective mutants (UbiB^D310N^, UbiB ^D310N^ and UbiB ^D310N^) were expressed in the Δ*ubiB*, Δ*ubiX*Δ*ubiB* and Δ*ubiX*Δ*ubiD* mutant strains. Strains containing an empty plasmid (vec) were used as controls. UQ_8_ content (ECD detection) of lipid extracts from Δ*ubiB* **(A)**, Δ*ubiX*Δ*ubiB* **(B)** and Δ*ubiX*Δ*ubiD* **(C)** mutant strains. Percentages referred to in the main text are indicated. **(D)** OHB (mass detection M-H, m/z 681.5) contents of lipid extracts from the Δ*ubiX*Δ*ubiD* strain. Values are shown with means ± SD (n=3). ****P < 0.0001, ***P < 0.001, **P < 0.01, *P < 0.05, ns, not significant by unpaired Student’s test comparing each mutant UbiB to its corresponding WT (lines) for panel A, and to vector (vec) for panels C and D. ND: not detected.

### Homologs of UbiB proteins from bacteria lacking *ubiX* and *ubiD* genes improve the efficiency of the alternative decarboxylation step in *E. coli*

To further explore the potential involvement of UbiB proteins in the alternative decarboxylation of OHB, we studied UbiB proteins from *Francisella novicida* (UbiB*_Fn_*) and *Xanthomonas campestris* (UbiB*_Xc_*), two organisms that lack UbiX and UbiD in their genomes (22, 23). UbiB*_Fn_* and UbiB*_Xc_*have 48% and 47% amino acid sequence identity with UbiB from *E. coli*, respectively (Fig. S4). We found that both proteins partially rescued UQ_8_ biosynthesis in *E. coli* Δ*ubiB* cells, namely to 45% (UbiB*_Fn_*) and 65% (UbiB*_Xc_*) of the level obtained with WT UbiB (Fig. 5A). Interestingly, in the Δ*ubiX*Δ*ubiB* strain, UbiB*_Xc_*and UbiB*_Fn_* yielded approximately two- and threefold higher UQ_8_ levels than WT UbiB, respectively (Fig. 5B). Mutation of the Asp residue corresponding to *E. coli* UbiB^D310^ (Fig. S4) only mildly impacted the activity of UbiB*_Fn_*^D307N^ and UbiB_Xc_ ^D313N^ in Δ*ubiB* cells (Fig. 5A), but abrogated function in the Δ*ubiX*Δ*ubiB* cells (Fig. 5B). These results indicate that the conserved aspartate residue (D310 in *E. coli*, D307 in *F. novicida* and D313 in *X. campestris*) is required for decarboxylase activity but is dispensable for UQ□ biosynthesis when the UbiX/UbiD system is functional.

Next, we expressed UbiB proteins in the Δ*ubiX*Δ*ubiD* mutant strain to evaluate their capacity to enhance its UQ_8_ content. Expression of *E. coli* UbiB modestly increased UQ_8_ levels by 17% compared to the empty vector (Fig. 5C), and significantly decreased OHB levels (Fig. 5D). In contrast, UbiB*_Xc_* and UbiB*_Fn_* increased the UQ_8_ content more robustly, by 97 and 135%, respectively, compared to the empty vector (Fig. 5C). Concomitantly, a strong decrease of OHB content was observed (Fig. 5D). As expected, expression of UbiB^D310N^, UbiB*_Fn_*^D307N^ and UbiB_Xc_ ^D313N^ in the Δ*ubiX*Δ*ubiD* mutant strain had no significant effect on the UQ_8_ content, and OHB levels remained elevated (Fig. 5C and Fig. 5D). Collectively, these results indicate that UbiB proteins exhibit a conserved capacity to enhance decarboxylation of OHB into OPP. Furthermore, the higher efficiency of UbiB*_Fn_* and UbiB*_Xc_* indicates that UbiB likely serves as the primary decarboxylase in UQ biosynthesis for *F. novicida* and *X. campestris*, which lack UbiX and UbiD in their genomes.

### Highly conserved UbiB active site residues are important for the alternative decarboxylation activity

We performed a Clustal Omega alignment of 4,011 UbiB sequences from representative bacterial strains of *Pseudomonadota* (dataset S1) and identified a conserved HxDxHxxN signature motif (Fig. 6A), which is absent from the eukaryotic Coq8 homologs. Among the most highly conserved residues are two histidines (H286 and H290), one aspartate (D288), and one asparagine (N293). In the AlphaFold model of *E.*L*coli* UbiB, these residues cluster in a region that appears to define the active site (Fig. 6B). Three highly conserved lysine residues (K70, K153, and K407) and one aspartate (D310, above-discussed and analyzed) are also present within the predicted active site pocket (Fig. 6A and Fig. 6B). To evaluate the contribution of these residues to UbiB function, namely the transfer of OPP to the Ubi-complex, as assessed in the Δ*ubiB* strain, and the decarboxylation of OHB into OPP, as assessed in the Δ*ubiX*Δ*ubiB* strain, we generated point substitutions in the *E. coli* protein, resulting in four mutant strains. UbiB^D310N^ as well as UbiB^K153A^, UbiB^H286L^, and UbiB^H290L^ complemented the UQ_8_ biosynthesis defect of the Δ*ubiB* mutant strain, yielding approximately 66%, 103%, 85%, and 101% of the UQ_8_ level obtained with the WT protein, respectively (Fig. 6C). The impact of these mutations was stronger in the Δ*ubiX*Δ*ubiB* strain, with UbiB^K153A^ and UbiB^H286L^ showing reduced UQ_8_ levels (∼46% and 70%, respectively), while UbiB^D310N^ and UbiB^H290L^ did not produce detectable UQ_8_ (Fig. 6D). UbiB^K70A^ complemented both mutant strains, while UbiB^K407A^ unexpectedly increased UQ_8_ levels by ∼241% in the Δ*ubiX*Δ*ubiB* mutant strain compared to the control (Fig. 6D). Finally, the D288N and N293L mutations resulted in a strong reduction in UQ_8_ content, regardless of the genetic background (Fig. 6C and Fig. 6D). Overall, these results indicate that H286, H290, D310, and, to a lesser extent, K153 primarily contribute to decarboxylation, whereas D288 and N293 are additionally required for OPP delivery to the Ubi-complex. (Fig. 6E).

**Figure 6:**
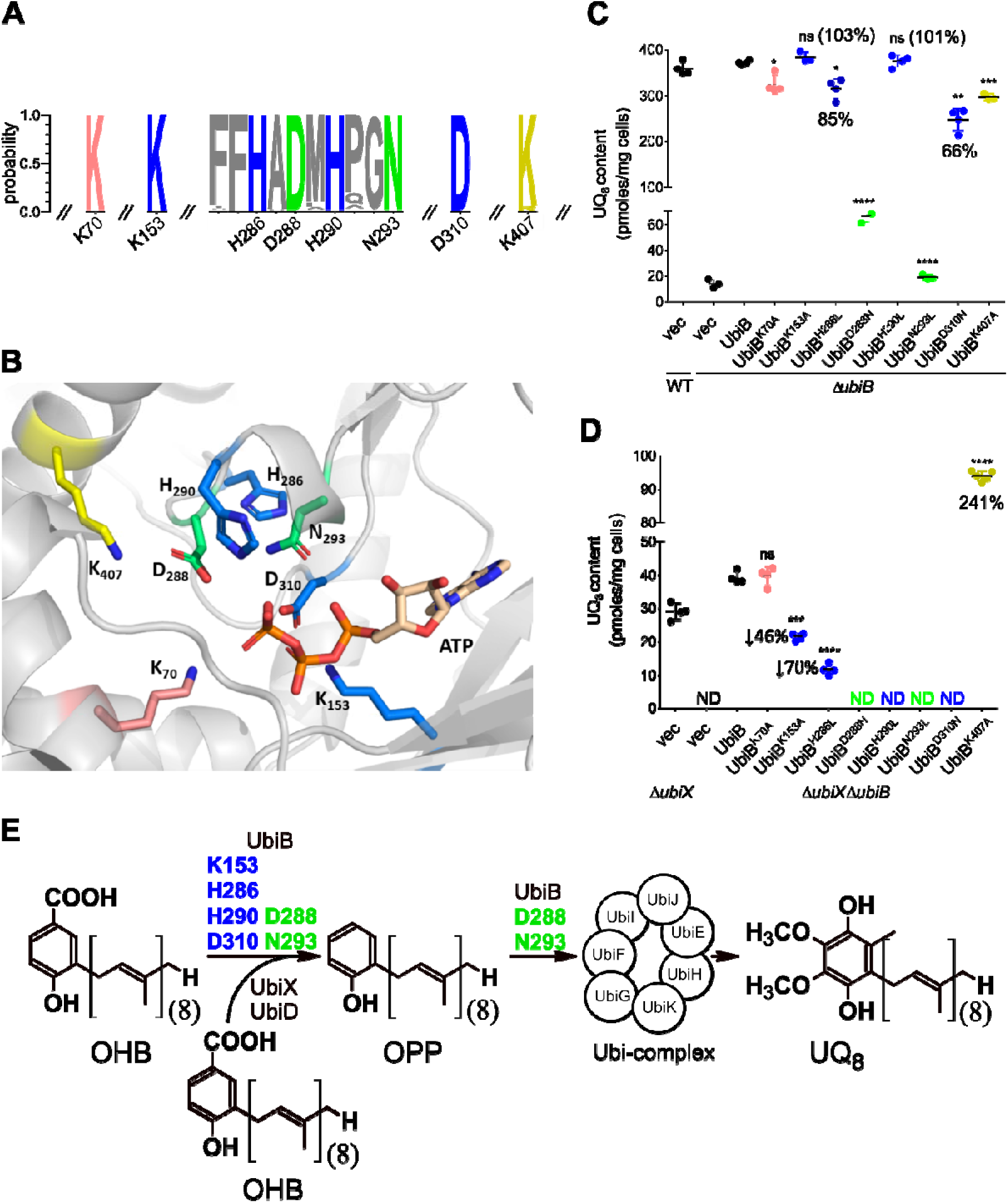
Mutational analysis of conserved residues localized in the predicted active site pocket of UbiB proteins. **(A)** Representation of the conservation of the signature sequence and selected residues over 4011 UbiB protein sequences as obtained with WebLogo3 (www.xeblogo.threeplusone.com). The numbering corresponds to the residues of the *E. coli* protein. **(B)** Three-dimensional model of the active site pocket of UbiB from *E. coli* generated with AlphaFold 3. An ATP molecule, shown in wheat, was docked into the active site using AlphaFold 3. Secondary structures of the model are represented in grey by PyMOL. Highly conserved residues are represented as sticks, with oxygen atoms in red and nitrogen atoms in dark blue. Backbone color is associated to the data shown in panels (C) and (D). UQ_8_ content in Δ*ubiB* **(C)** or Δ*ubiX*Δ*ubiB* **(D)** mutated strains transformed with plasmids expressing UbiB variants. Percentages referred to in the main text are indicated. In pink, mutation with minor or without effect in the two genetic backgrounds; in yellow, mutation which increases UQ_8_ content only in Δ*ubiX*Δ*ubiB* strain; in blue, mutations which cause a strong decrease of the UQ_8_ content mainly in Δ*ubiX*Δ*ubiB* strain; in green, mutations which cause a strong decrease of the UQ_8_ content in the two genetic backgrounds. Values are shown with means ± SD (n=3). ****P < 0.0001, ***P < 0.001, **P < 0.01, *P < 0.05, ns, not significant by unpaired Student’s test comparing to WT UbiB. ND: not detected. **(E)** The proposed functions of the UbiB protein in the UQ biosynthetic pathway and the amino acids suggested as important for each function. Abbreviations: OHB, octaprenyl-hydroxybenzoic acid; OPP, octaprenylphenol; UQ_8_, ubiquinone 8. ND: not detected.

### The alternative decarboxylase activity is linked to UbiB ATPase activity

As previously demonstrated in the literature, the UbiB protein family exhibits an evolutionarily conserved ATPase activity, from *E. coli* to humans (18). Interestingly, all conserved residues of the signature motif are located in close proximity to the ATP molecule docked in the predicted active-site pocket of UbiB. The ATP-binding mode predicted by AlphaFold 3 is comparable to that of AMP-PNP (adenylyl imidodiphosphate, a non-hydrolysable analogue of ATP) observed in the structure of human Coq8 (PDB: 5I35), the mammalian homolog of UbiB (Fig. 6B). To investigate the connection between ATPase and decarboxylase activities, we purified a maltose-binding protein (MBP)-His_8_-tagged version of *E. coli* UbiB lacking its predicted C-terminal transmembrane domains (UbiB^ΔC47^) (Fig. 7A).

**Figure 7:**
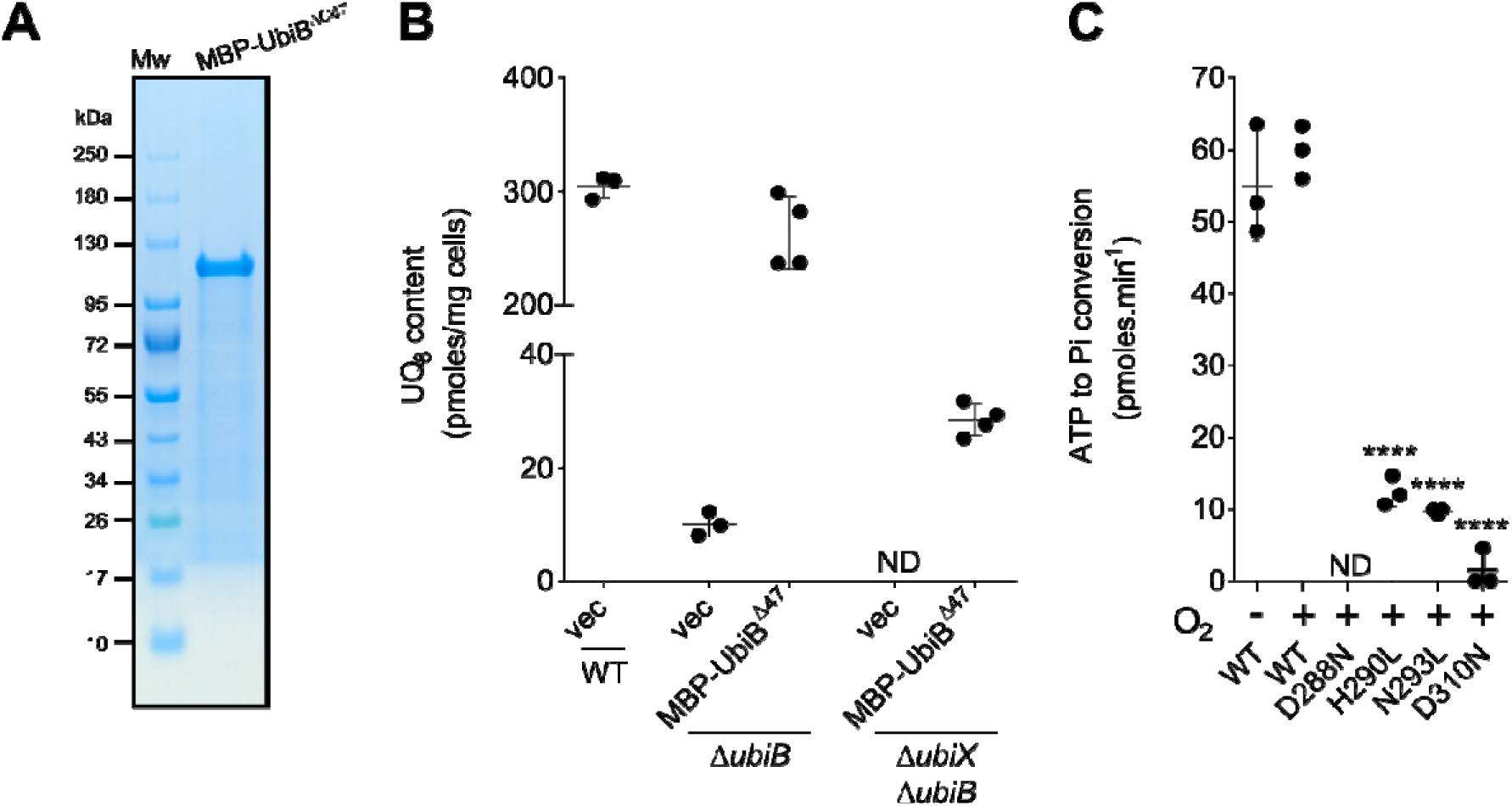
Identification of conserved residues required for UbiB ATPase Activity. **(A)** Coomassie Blue-stained SDS-PAGE analysis of purified MBP-UbiB^ΔC47^. Mw: molecular weight marker. **(B)** UQ_8_ content of lipid extracts from Δ*ubiB* and Δ*ubiX*Δ*ubiB* mutated strains containing either an empty plasmid (vec) or a plasmid expressing MBP-UbiB^ΔC47^. Values are shown with means ± SD (n=3 to 4). **(C)** Malachite green ATPase assay of 0.7 µM MBP-UbiB^ΔC47^ and its point mutants (D288N, H290L, N293L and D310N). ****P < 0.0001 by unpaired Student’s test comparing to WT UbiB. ND: not detected.

Notably, MBP-UbiB^ΔC47^ was active *in vivo* in the Δ*ubiB* and Δ*ubiX*Δ*ubiB* strains (Fig. 7B), showing that neither the MBP-tag nor the C-terminal truncation interfered with UbiB’s activities. The ATPase activity of MBP-UbiB^ΔC47^ and its mutants was assessed with the malachite green phosphate assay kit using a previously described reaction mixture containing 2-propyl-phenol as an activator (18). We also generated the D288N, H290L, N293L, and D310N variants, all of which were expressed at similar levels (Fig. S5). We demonstrated that the in vitro ATPase activity was independent of oxygen and was reduced by approximately 79%, 84%, and 97% respectively in the H290L, N293L, and D310N mutants, whereas it was abolished in the D288N mutant (Fig. 7C). These results are consistent with those obtained in the *in vivo* decarboxylation assay (Fig. 6D), and support the conclusion that ATPase activity is critical for UbiB-mediated decarboxylation. In contrast, the ATPase activity appears dispensable for the transfer of OPP to the Ubi-complex, as confirmed by the high *in vivo* activity of the H290L and D310N mutants in Δ*ubiB* cells (Fig. 6C).

## Discussion

UbiB proteins belong to an atypical protein kinase-like superfamily that is broadly conserved across prokaryotic and eukaryotic lineages. UbiB proteins play an essential role in UQ biosynthesis (17, 18, 30), but their exact function remains mysterious. Their intrinsic ATPase activity, a conserved biochemical feature across orthologs from *E. coli* to humans, is enhanced by phenolic compounds structurally analogous to intermediates of the UQ pathway (18). Based on these observations, it has been proposed that UbiB proteins, owing to their membrane localization, mediate ATP-driven extraction of hydrophobic UQ precursors from the membrane environment, thereby facilitating their transfer to soluble or membrane-associated enzymes involved in UQ biosynthesis (3). Beyond this role, UbiB proteins have also been proposed to regulate the intracellular distribution of UQ in eukaryotic cells (30). Here, we provide evidence that several bacterial UbiB proteins are able to decarboxylate OHB into OPP *in vivo* (Fig. 6E). This activity was most clearly observed in *E. coli* in the absence of UbiX/UbiD, the main decarboxylase system. The alternative decarboxylation activity was found to be O_2_-dependent and linked to UbiB ATPase activity. Consequently, this discovery challenges the current model of UbiB function and calls for a reassessment of the role of its ATPase activity in UQ biosynthesis, which are discussed below.

The UQ biosynthesis pathway has been extensively studied in *E. coli*, but our recent work demonstrated an unsuspected diversity of this pathway across bacteria within the phylum *Pseudomonadota*, particularly regarding the hydroxylation steps (19, 20, 24). Computational and experimental analyses presented in this study extend this observation to the decarboxylation step. Indeed, although the well-characterized UbiX/UbiD decarboxylation system is widespread among UQ-synthesizing bacteria, about one-third of the analyzed *Pseudomonadota* genomes lack homologs of UbiX and UbiD. This contrasts with other ubiquitous Ubi-proteins, namely UbiA, UbiB, UbiE, and UbiG, which are consistently present (24). Thus, alternative systems to UbiX and UbiD likely catalyze the decarboxylation step in a substantial fraction of *Pseudomonadota*, as previously hypothesized for *F. novicida* and *X. campestris* bacteria (22, 23). Recently, a member of the class A FMO, was shown to be involved in the decarboxylative hydroxylation step of the UQ biosynthesis pathway in *R. capsulatus* (21). Interestingly, in this organism, this FMO operates in parallel with the UbiX/UbiD system, enabling the decarboxylation step to proceed under aerobic and anaerobic conditions, respectively. The phylogenetic distribution of this enzyme is unknown, but it is unlikely that it represents the alternative decarboxylase system in all species lacking the UbiX/UbiD system, because neither *E. coli*, *F. novicida*, nor *X. campestris* harbor homologs in their genomes. However, we note that *E. coli* and *R. capsulatus* share an oxygen-dependent decarboxylase system. Moreover, we showed that most of the UQ-producing bacteria lacking both UbiD and UbiX rely primarily on O_2_-dependent respiration. This observation is consistent with our biochemical findings and supports the idea that oxygen plays a central role in the activity of alternative decarboxylases that compensate for the absence of the UbiX/UbiD system.

UbiB is a central component of the UQ biosynthesis pathway. The marked accumulation of OPP in a Δ*ubiB* strain indicates an early block in the pathway, consistent with the proposed primary role of UbiB in the extraction of early intermediates from the membrane for transfer to the soluble Ubi-complex (3). Our results demonstrate that UbiB is also essential in *E. coli* to sustain OHB decarboxylation when the UbiX/UbiD system is disabled. Moreover, mutants such as D310N and H290L, which impair decarboxylation without affecting canonical UbiB function, provide strong evidence that UbiB can also function as a decarboxylase. We further demonstrate the superior efficiency of UbiB proteins from *F. novicida* and *X. campestris* compared to endogenous *E. coli* UbiB in decarboxylating OHB. These bacteria may therefore dispense with an additional decarboxylase system such as that of UbiX/UbiD. An important question to address is why this activity has been conserved in UbiB from *E. coli*, despite the presence of the UbiX/UbiD system. One plausible explanation is that this secondary activity serves as a safeguard mechanism, ensuring that residual OHB molecules escaping the UbiX/UbiD system are decarboxylated before incorporation into the Ubi-complex.

Alignment of UbiB sequences from representative *Pseudomonadota* strains revealed a highly conserved motif, whose residues co-localize within the predicted active site of the *E. coli* UbiB AlphaFold model. These residues are located near the modeled ATP molecule. As mutations in several conserved residues primarily impair the *in vivo* UbiB-dependent decarboxylase activity, as well as the *in vitro* ATPase activity of UbiB, we propose that ATPase activity is critical for UbiB function, at least for catalyzing the decarboxylation reaction. However, residues D288 and N293 are also important for the canonical activity of UbiB, suggesting that all activities of UbiB are likely interconnected. The mechanism underlying the ATP-dependent decarboxylation activity of UbiB remains unknown and our attempts to develop an *in vitro* decarboxylation assay from 3-farnesyl-4-hydroxybenzoic acid have so far been unsuccessful. Several explanations may account for this negative result. The solubilization step using DDM (*n*-dodecyl-β-D-maltoside) may have partially destabilized the protein, potentially impairing its ability to bind its substrate. Another possibility is that UbiB requires interaction with the Ubi-complex for decarboxylation to occur. Regarding the mechanism, it may resemble that of mevalonate diphosphate decarboxylase, which possesses an ATP-dependent decarboxylase activity (31). We have also demonstrated that the *in vivo* decarboxylase activity of UbiB relies on O_2_; however, the role played by O_2_ remains enigmatic. As the three hydroxyl groups added to the phenyl ring of UQ biosynthetic intermediates come from the Ubi-FMO (32), we suppose that O_2_ is unlikely to be incorporated into the decarboxylation product generated by UbiB. This hypothesis is consistent with the mechanism proposed for HemF in *E. coli*. Indeed, this enzyme, which catalyzes an oxidative decarboxylation step in the porphyrin biosynthesis pathway, uses molecular oxygen only as a terminal electron acceptor and produces hydrogen peroxide as a by-product (33).

Collectively, our data assign a new role to UbiB proteins as decarboxylases. Given that UbiB is ubiquitous in UQ biosynthesis across *Pseudomonadota* (24), it will be interesting to investigate whether UbiB ancestrally possessed decarboxylase activity.

## Material and methods

### Bacterial strains and plasmid constructions

*E. coli* strains, plasmids and primers used in this study are respectively listed in Tables S2, S3 and S4 in the supplemental material. The Δ*ubiX*Δ*ubiB* double knock-out (KO) strain was constructed by P1 transduction using MG1655 Δ*ubiX* as the recipient strain and MG1655 Δ*ubiB*::cat as the donor strain using an established protocol (https://barricklab.org/twiki/bin/view/Lab/ProtocolsP1Transduction). The presence of the *ubiB*::cat mutation was confirmed by colony polymerase chain reaction (PCR) with primers flanking the *ubiB* locus.

The Δ*ubiX*Δ*ubiD* double KO strain has been kindly provided by Dr Laurent Loiseau (LCB in Marseille). The two single KO strains, Δ*ubiX* and Δ*ubiD* were cured with pCP20 to respectively yield Δ*ubiXc* and Δ*ubiDc*. *E. coli ubiB* gene (*ubiB*) was obtained by PCR amplification using the MG1655 genome as the template and the oligonucleotides 5’-*ubiB_Ec_* and 3’-*ubiB_Ec_* listed in Table S4. Insert was EcoRI/XbaI digested and inserted into EcoRI/XbaI-digested pTrc99a plasmid. *X. campestris ubiB* gene (*ubiB_Xc_*) was synthesized by the “Genecust” company and cloned into pTrc99a vector downstream of lactose-inducible promotor using EcoRI (5′ end) and BamHI (3′ end) restriction enzymes. The nucleotide sequence was optimized for expression in *E. coli* and is available in supplementary Dataset S2. Two single mutant *ubiB* (K70A) and *ubiB* (K407A) genes were synthesized by the “Genecust” company and cloned into pTrc99a vector downstream of lactose-inducible promotor using EcoRI (5′ end) and XbaI (3′ end) restriction enzymes. Point mutations of *ubiB* (K153A, H286L, D288N, H286L, H290L and D310N), of *ubiB_Xc_* (D313N) and of *ubiB*_Fn_ (D307N) genes were introduced by PCR-based site-directed mutagenesis (Table S4). The gene encoding the MBP-His_10_-tagged version of *E. coli* UbiB, lacking its predicted C-terminal transmembrane domains (UbiB^ΔC47^), was constructed as follows. The gene encoding UbiB^ΔC47^was amplified by PCR using pTrc99a-*ubiB* as a template and primers listed in Table S4, then cloned into the pETM40 vector using SLIC (Sequence and Ligation-Independent Cloning), yielding the pETM40-MBP-UbiB^ΔC47^ vector. Point mutations (D288N, H290L, N293L and D310N) were introduced by PCR-based site-directed mutagenesis (Table S4). All genetic constructs used in this study were verified by DNA sequencing (Eurofins).

### Media and growth conditions

Bacterial strains were cultured in lysogeny broth (LB) medium. When required, ampicillin (100 μg/mL) and chloramphenicol (35 μg/mL) were added from aqueous stock solutions sterilized by filtration through 0.2-μm filters. For aerobic cultures, strains were grown in glass tubes (15 cm long, 2 cm in diameter) at 37 °C with shaking at 180 rpm. Fresh medium (5 mL) was inoculated with 100 μL of an overnight (O.N.) culture and incubated O.N. For anaerobic cultures, Hungate tubes were used as previously described (4) The LB medium was supplemented with 2.5 mg/L resazurin as an anaerobic indicator, 100 mg/L L-cysteine (pH adjusted to 6 with NaOH) to scavenge residual oxygen, and anti-foam (Sigma Life Science, 0.5 mL/L). The medium (13 mL) was distributed into Hungate tubes and deoxygenated by bubbling high-purity argon for 30 min. The tubes were then sealed and autoclaved. Precultures were grown O.N. at 37 °C in Eppendorf tubes filled to the top with LB medium. Hungate tubes were subsequently inoculated through the septum using disposable syringes and needles with 100 μL of preculture and incubated at 37 °C without agitation. Antibiotics were added post-autoclaving, simultaneously with the preculture inoculation. For *in vivo* complementation assays, Δ*ubiB*, Δ*ubiX*Δ*ubiB*, and Δ*ubiX*Δ*ubiD* mutant strains from *E. coli* were transformed with either empty pTrc99a vectors or pTrc99a carrying the *ubiB* genes and their mutants. Transformants were selected on LB agar supplemented with ampicillin (100 μg/mL), and individual clones were used to inoculate O.N. cultures. For the anaerobic-to-aerobic shift assay, the Δ*ubiDc* strain was first precultured anaerobically in Hungate tubes for ∼10 hours. A 500-mL bottle with a two-port cap, fitted with plastic tubing for continuous argon injection to maintain anaerobiosis, was filled with 250 mL of pre-deoxygenated LB medium. The culture was inoculated at an OD_600_ of 0.1 and incubated at 37 °C under high-purity argon bubbling for ∼2 hours until reaching an OD_600_ of 0.35. At this point, 500 μL of chloramphenicol (200 μg/mL) was injected. After 30 minutes, the bottles were unsealed, and a 5 mL sample was collected for lipid extraction and quinone analysis. The remaining culture was incubated at 37 °C with shaking at 180 rpm. Samples (5 mL) were collected at 15, 30, 45, 60, 90, 120, and 180 minutes, as well as 4 and 24 hours after exposure to ambient air, for lipid extraction and quinone content analysis.

### Lipid extractions and quinone analysis

First, 5 mL of each culture was cooled on ice for at least 30 min before centrifugation at 3,200 g at 4 °C for 10 min. Cell pellets were washed in 1 mL ice-cold phosphate-buffer saline and transferred to preweighted 1.5 mL Eppendorf tubes. After centrifugation at 12,000 g at 4 °C for 1 min, the supernatant was discarded, the cell wet weight was determined and pellets were stored at −20 °C until lipid extraction, if necessary. Quinone extraction from cell pellets was performed as previously described (10). The dried lipid extracts were resuspended in 100 µL ethanol, and a volume corresponding to 1 mg of cell wet weight was analyzed by HPLC electrochemical detection-mass spectrometry (ECD-MS) with a BetaBasic-18 column at a flow rate of 1 mL/min with a mobile phase composed of 50% methanol, 40% ethanol, and 10% of a mix (90% isopropanol, 10% ammonium acetate (1 M), and 0.1% formic acid). MS detection was performed with a MSQ spectrometer (Thermo Scientific) with electrospray ionization in positive (probe temperature, 400°C; cone voltage, 80 V) and negative mode (probe temperature, 450°C; cone voltage, 60 V). Single-ion monitoring detected the following compounds: UQ_8_ (M+NH_4_^+^), m/z 744-745, 6–10 min, scan time of 0.2 s; UQ_10_ (M+NH_4_^+^), m/z 880–881, 10-17 min, scan time of 0.2 s; OPP (M+NH_4_ ^+^), 656-657, 5-9 min, scan time of 0.4 s; and OHB (M-H), 681-682, 6-10 min, scan time of 0.2 s. MS spectra were recorded between m/z 600 and 900 with a scan time of 0.3 s. ECD and MS peak areas were corrected for sample loss during extraction on the basis of the recovery of the UQ_10_ internal standard and then were normalized to cell wet weight. The peaks of UQ_8_ and the biosynthetic intermediates obtained with electrochemical detection or MS detection were quantified with a standard curve of UQ_10_.

### Expression and purification of MBP-UbiB**^Δ^**^C47^ from *E. coli*

The overexpression of *E. coli* MBP-UbiB^ΔC47^ was carried out in chemically competent *E. coli* Lemo21(DE3) cells (New England Biolabs), transformed with the pETM40-MBP-UbiB^ΔC47^ plasmid. A starter culture of 100 mL LB medium supplemented with 50 µg/mL kanamycin was grown overnight and used to inoculate a 6 L large-scale culture. Cells were incubated at 37 °C until the optical density at 600 nm (OD_600_) reached 0.4. Protein expression was induced by adding isopropyl-1-thio-β-D-galactopyranoside (IPTG) to a final concentration of 0.4 mM. Cultures were then incubated for 16 h at 16 °C before harvesting cells by centrifugation at 9,000 × g for 10 min at 4 °C. Cell pellets were stored at −80 °C until further use. Cells were resuspended in 10 mM PBS buffer (pH 7.4) containing 500 mM NaCl, 5% (v/v) glycerol, 1 mM DTT, 1 mM PMSF, and 0.5% (w/v) n-dodecyl-β-D-maltoside (DDM). Lysis was performed by discontinuous sonication for 15 minutes on ice. The lysate was centrifuged at 193,000 × g for 45 minutes to obtain the soluble fraction of MBP-UbiB^ΔC47^. The supernatant was loaded onto a 5 mL Hitrap Excel™ column (Cytiva) pre-equilibrated with buffer containing 10 mM PBS (pH 7.4), 200 mM NaCl, 5% (v/v) glycerol, 1 mM DTT, 1 mM PMSF, and 0.5% (w/v) DDM. Bound proteins were eluted using a linear imidazole gradient (0–500 mM) in buffer A (50 mM Tris-HCl pH 7.5, 300 mM NaCl, 5% glycerol, 1 mM DTT, and up to 500 mM imidazole). MBP-UbiB^ΔC47^ eluted at approximately 70 mM imidazole. A total of 30 mg of the eluted protein were subjected to size-exclusion chromatography using a HiLoad 16/600 Superdex™ 200 pg column, equilibrated in buffer B (10 mM PBS pH 7.4, 200 mM NaCl, 5% (v/v) glycerol, 1 mM DTT, 1 mM PMSF). The main peak was collected and further purified on an MBPTrap HP™ column (5 mL, Cytiva), pre-equilibrated with buffer B. Elution was achieved using a step of 10 mM maltose in the same buffer. Protein concentration was determined using the Bradford assay (Bio-Rad), with BSA as the standard. Pure protein fractions were pooled, concentrated to 5 mg/mL, flash-frozen in liquid nitrogen, and stored at −80 °C.

### ATPase assays

ATPase assays were performed using malachite green phosphate assay kit (Sigma Adrich) according to the manufacturer’s instruction with the following modifications. In a typical ATPase assay, all solutions were diluted in HBS (20 mM HEPES buffer at pH 7.5, 150 mM NaCl). ATP (100 µM final), MgCl_2_ (4 mM final), 2-propylphenol (1 mM final), COQ 1 (0,5 mM final), reduced Triton X-100 (1 mM final), MBP-UbiB^ΔC47^ (0.7 µM final). Reactions were incubated at room temperature for 15 min and 20µL of reagent was added and incubated (room temperature, 15 min). Absorbance was measured at 620 nm. The standard curve ranged from 0-50 µM phosphate.

#### Sequences and alignment

A total of 19,777 Pseudomonadota genomes were retrieved from the NCBI database in August 2023 (NCBI). Redundant entries were removed by selecting one representative genome per species, resulting in a dataset of 4,107 genomes as in (24). Gene annotations for UQ and menaquinone biosynthesis through the Men biosynthetic pathway were derived from a previous study where hits were obtained with hmmscan from the HMMER suite (v3.3.2) using a set of previously described HMM protein profiles (24). Bioinformatics analyses were performed on a dataset of 3,928 *Pseudomonadota* genomes producing UQ, selected based on the presence of core UQ biosynthesis genes, i.e., *ubiA*, *ubiB*, *ubiE*, and *ubiG*. Gene annotations for UQ biosynthesis and identification of the Men pathway for menaquinone biosynthesis were derived from a prior study (24). For sequence analysis, 4,011 UbiB protein sequences were aligned using Clustal Omega (v1.2.4) with default parameters (34) and representation of the conservation of selected residues was performed using Jalview 2.11.5.1.

### Metabolism annotation

To obtain metabolic data, we utilized a table from a study that compiled information from multiple sources (35). As in a previous study (24), we simplified the categories in the table by grouping species annotated as “obligate aerobic” and “aerobic” into an “aerobic” category, and those labeled as “obligate anaerobic” and “anaerobic” into an “anaerobic” category. Metabolic information was successfully retrieved for 1,037 species in our dataset.

## Data availability

The data underlying this article are provided as part of the main text and the supplementary data.

## Author contributions

L.P., M.L., M.F. and F.P. designed the research and L.P. and M.L. obtained the fundings. S-C.C. and S.S.A. performed all the bioinformatic analyses. K.K., B.F., J.M., T.A.D., A.L. and O.L. performed the experimental analyses and designed the corresponding figures. S-C.C. designed all other figures. L.P. and M.L. wrote the original version of this manuscript with contributions from all co-authors. All authors agree with this version of the manuscript.

## Acknowledgments

This work was supported by the French National Research Agency and by the Grenoble Alpes University through the grants from the Agence Nationale de la Recherche, ANR-21-CE02-0018 (S.C. and S.S.A), ANR-23-CE44-0015 (M.L.) and ANR-25-CE11-3606 (L.P.). The PhD of T.A.D was supported by a grant of the Graduate Schools/EUR from Grenoble Alpes University (ANR-17-EURE-0003).

