## Supplementary material for "UbiB proteins mediate an ATP-dependent decarboxylation step in bacterial ubiquinone biosynthesis": Fig.S1 to S5, Tables S2 to S4, Dataset S2

France

^3^Present address: Univ. Grenoble Alpes, CNRS, INRAE, IRD, Grenoble INP, IGE, 38000 Grenoble, France

Fig. S1: Mass spectrum (M-H) and chemical structure of OHB accumulated in the lipid extracts of the Δ*ubiX*Δ*ubiD* mutant strain.

Fig. S2: UQ_8_ content in the *E. coli* Δ*ubiXc* and Δ*ubiDc* single mutant strains.

Fig. S3: Restoration of UQ_8_ biosynthesis in *E. coli* Δ*ubiDc* mutants following an anoxic-to-oxic shift.

Fig. S4: Alignment of UbiB proteins.

Fig. S5: SDS-PAGE analysis of the different purified MBP-UbiB^ΔC47^ variants.

Table S2. Strains used in this study.

Table S3. Plasmids used in this study.

Table S4. Primers used in this study.

Dataset S2. Optimized sequence of *X. campestris* *ubiB* gene.

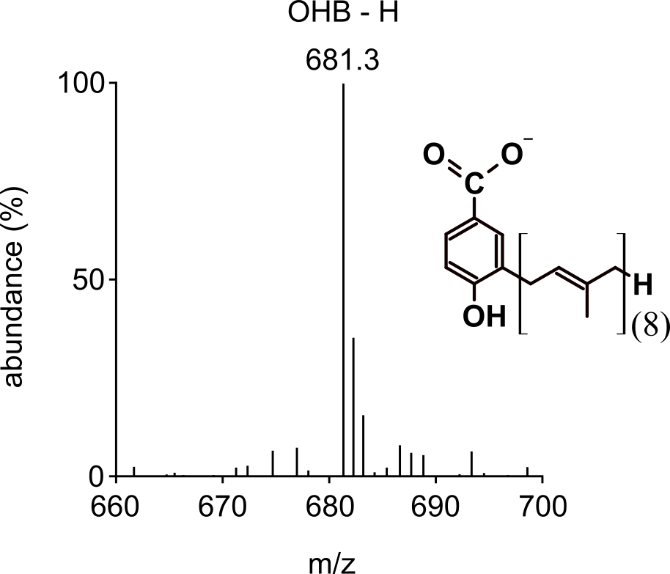

**Figure S1: Mass spectrum (M-H) and chemical structure of OHB accumulated in the lipid extracts of the Δ*ubiX*Δ*ubiD* mutant strain.**

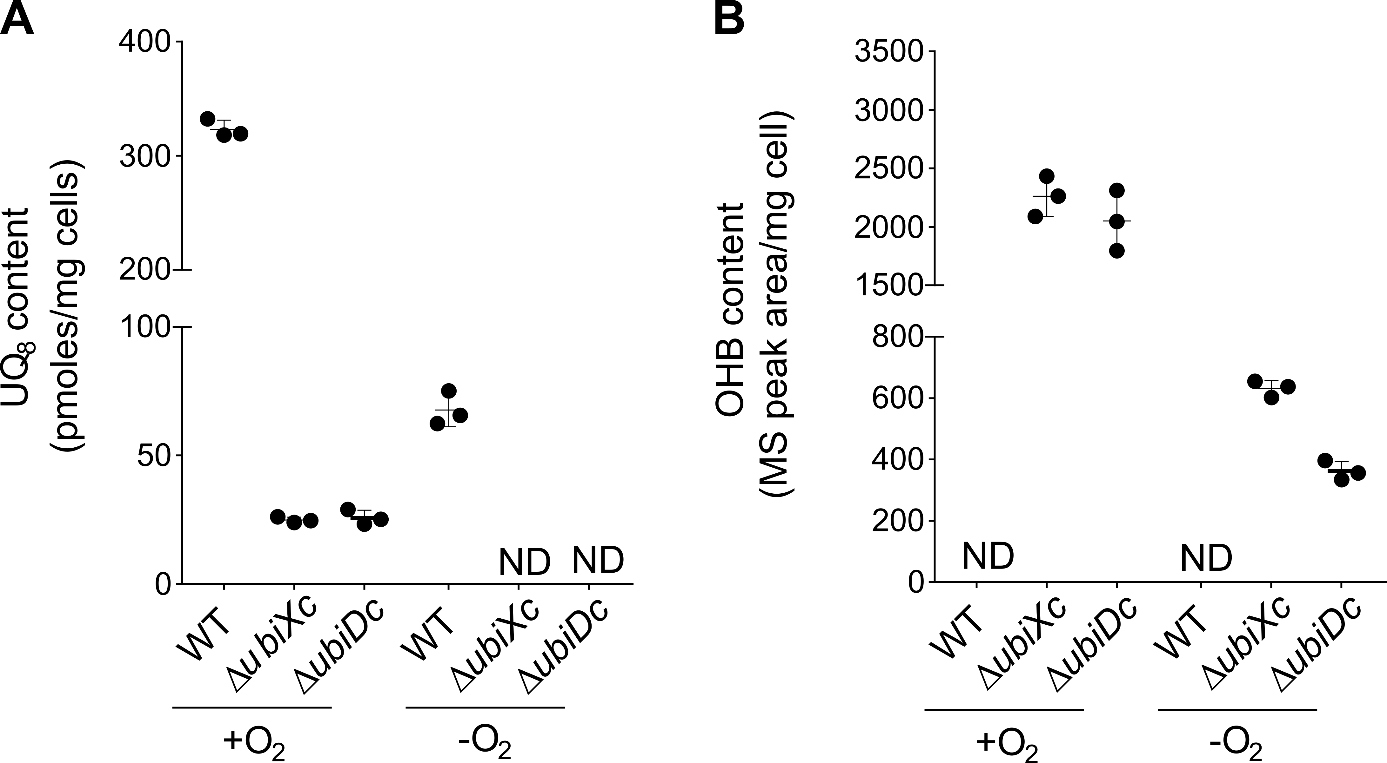

**Figure S2:** **UQ_8_ content in the *E. coli*** Δ***ubiXc* and** Δ***ubiDc* single mutant strains.** Analysis of lipid extracts from 1 mg of *E. coli* MG1655 (WT), Δ*ubiXc* or Δ*ubiDc* cells grown aerobically (+O_2_) or anaerobically (-O_2_) in LB medium. **(A)** UQ_8_ content (ECD detection) in cells. **(B)** OHB content (MS peak aera, mass detection M-H with m/z 681.3) in cells. Values are means ± SD (n=3). ND: not detected.

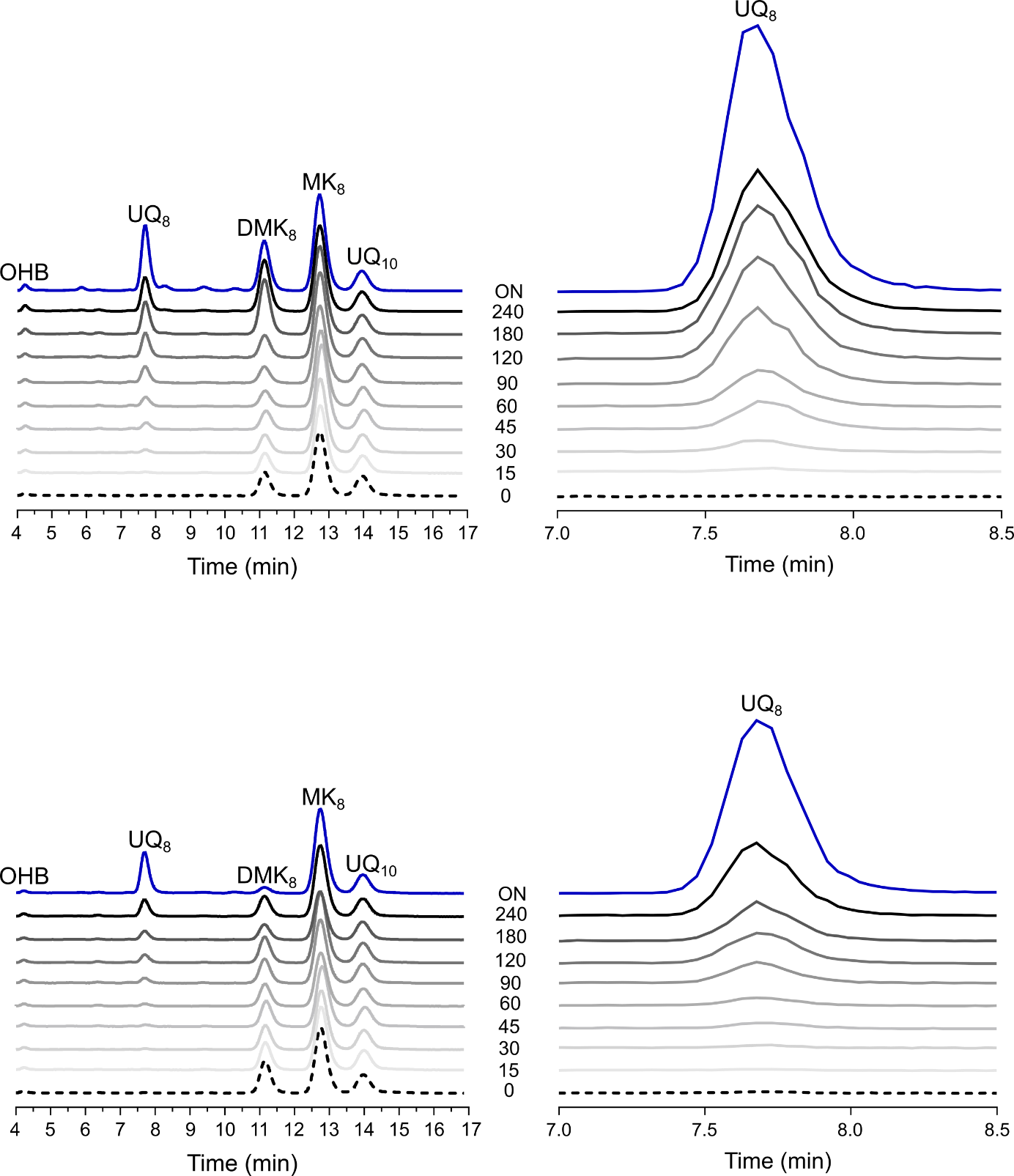

**Figure S3:** **Restoration of UQ_8_ biosynthesis in *E. coli* Δ*ubiDc* mutants following an anoxic-to-oxic shift.** UQ_8_ content was followed for 240 min and then after over-night (ON) culture in the bacteria treated without **(A and B)** or with **(C and D)** chloramphenicol. **(A and C)** HPLC-ECD analysis of lipid extracts from 1 mg of cells after growth in LB medium. The peaks corresponding to OHB, UQ_8_, DMK_8_, MK_8_ and the UQ_10_, as a standard, are indicated. **(B and D)** MS peaks (mass detection M+NH_4_^+^, m/z 744.5) used to quantified UQ_8_ content in cells. Chromatograms are representative of results from two independent experiments. Abbreviations: OHB, octaprenyl-hydroxybenzoic acid; UQ_8_, ubiquinone 8; DMK_8_, demethylmenaquinone; MK_8_, methylmenaquinone and UQ_10_, ubiquinone 10.

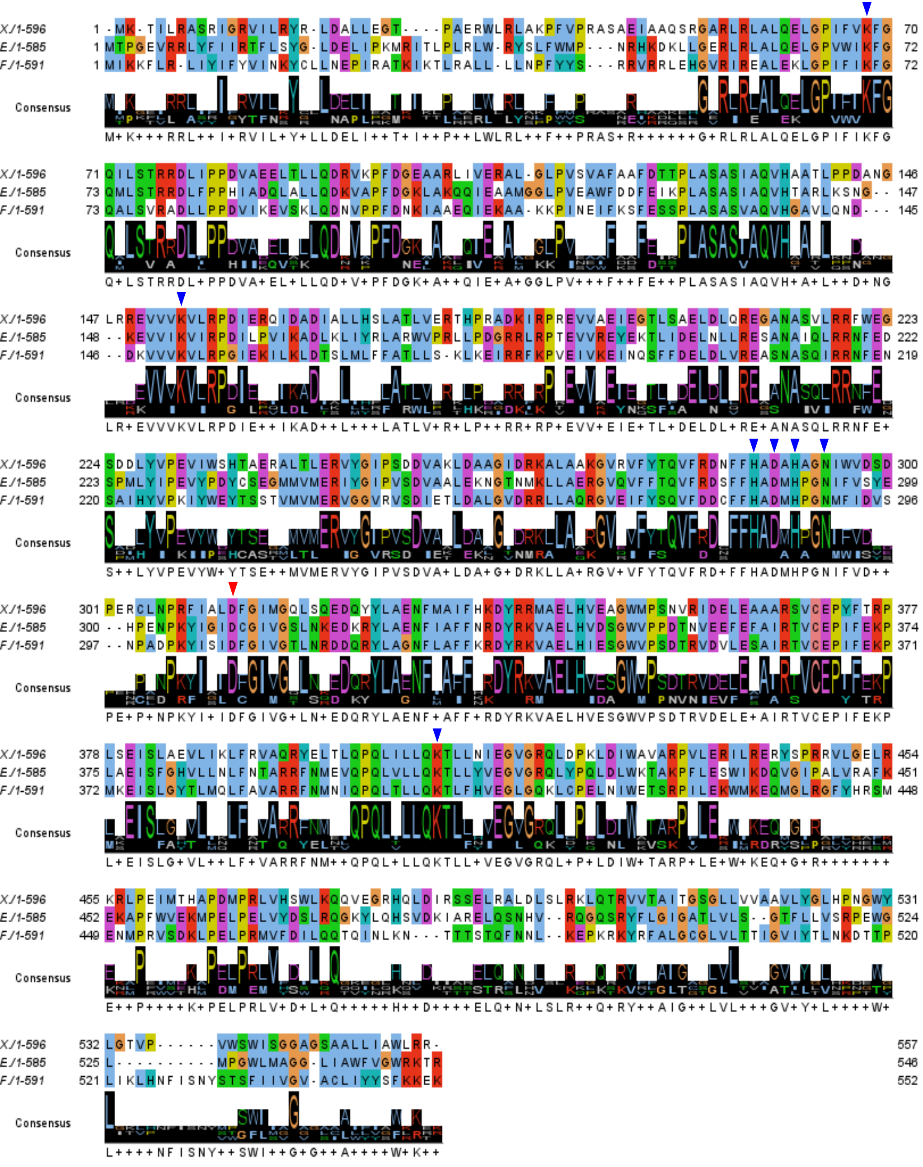

**Figure S4:** **Alignment of UbiB proteins.** Multiple sequence alignment of UbiB proteins from *E. coli* (*E*), *X. campestris* (*X*) and *F. novicida* (*F*) generated using ClustalW2 and analyzed with Jalview v2.0.1. The conserved aspartate residue, i.e., D310, D313 and D307 of UbiB proteins from *E. coli*, *X. campestris* and *F. novicida*, respectively, is indicated by a red triangle above the sequences. The blue triangles correspond to conserved residues within the predicted active site pocket of UbiB proteins (see Fig. 6).

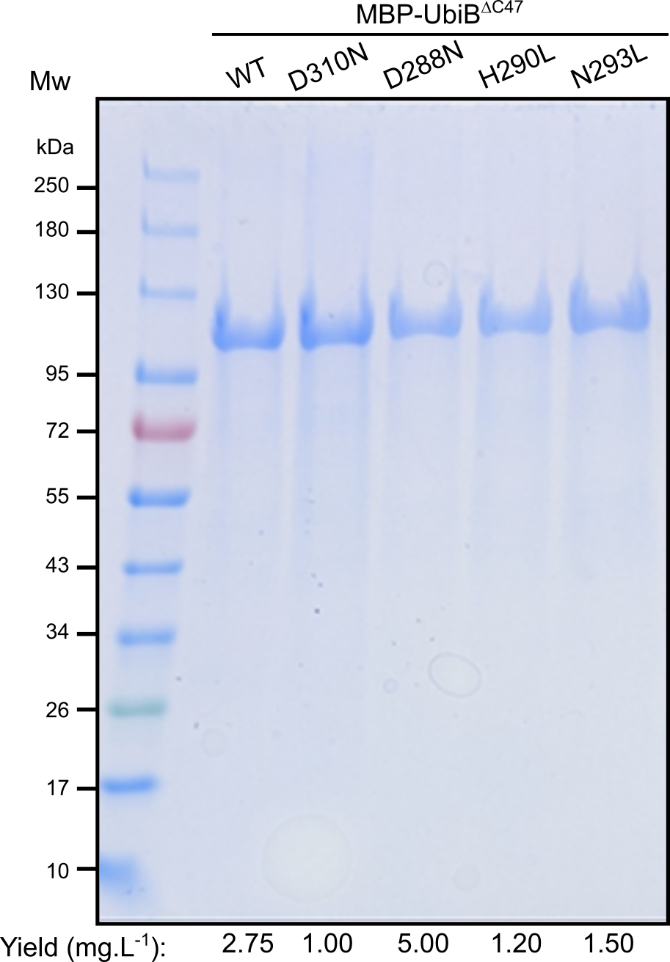

**Figure S5: SDS-PAGE analysis of the different purified MBP-UbiB^ΔC47^ variants.** D310N, D288N, N293L and H290L variants were obtained following a three-step purification procedure as described in the Materials and Methods section. The yield of each purified protein is indicated (mg·L⁻¹). For each sample, 10 µg of each protein were loaded onto the gel. Proteins were visualized by Coomassie Brilliant Blue staining.

| **Strain** | **Genotype** | **Construction** | **Source** |
| --- | --- | --- | --- |
| MG1655 | Wild type |  | Lab strain |
| ∆*ubiB* | MG1655 *ubiB*::cat |  | (1) |
| ∆*ubiDc* | as MG1655 but ∆*ubiD* | ∆*ubiD* cured with pCP20 | This study |
| ∆*ubiXc* | as MG1655 but ∆*ubiX* | ∆*ubiX* cured with pCP20 | This study |
| ∆*ubiX*∆*ubiB* | ∆*ubiX*∆*ubiB*::cat | ∆*ubiXc* + P1/*ubiB*::cat | This study |
| ∆*ubiX*∆*ubiD* | ∆*ubiX*∆*ubiD*::cat |  | kindly provided by Dr Laurent Loiseau |

**Table S2. Strains used in this study.**

**Table S3. Plasmids used in this study.**

*Ec*: *E. coli*, *Fn*: *F. novicida*, *Xc*: *X. campestris*.

| **plasmid** | **Description** | **source** |
| --- | --- | --- |
| pTrc99a | Vector for inducible gene expression (Amp^R^) | Ref 306986 (pubmed) |
| pTrc99a-*ubiB* | pTrc99a carrying *ubiB_Ec_* (EcoRI/XbaI) | This study |
| pTrc99a-*ubiB_Fn_* | pTrc99a carrying *ubiB_Fn_* (EcoRI/BamHI) | (2) |
| pTrc99a-*ubiB_Xc_* | pTrc99a carrying *ubiB_Xc_* (EcoRI/BamHI) | This study |
| pTrc99a-*ubiB* (D310N) | pTrc99a carrying *ubiB_Ec_* (D310N) (EcoRI/XbaI) | This study |
| pTrc99a-*ubiB* (D288N) | pTrc99a carrying *ubiB_Ec_* (D288N) (EcoRI/XbaI) | This study |
| pTrc99a-*ubiB* (H286L) | pTrc99a carrying *ubiB_Ec_* (H286L) (EcoRI/XbaI) | This study |
| pTrc99a-*ubiB* (H290L) | pTrc99a carrying *ubiB_Ec_* (H290L) (EcoRI/XbaI) | This study |
| pTrc99a-*ubiB* (N293L) | pTrc99a carrying *ubiB_Ec_* (N293L) (EcoRI/XbaI) | This study |
| pTrc99a-*ubiB* (K153A) | pTrc99a carrying *ubiB_Ec_* (K153A) (EcoRI/XbaI) | This study |
| pTrc99a-*ubiB* (K70A) | pTrc99a carrying *ubiB_Ec_* (K70A) (EcoRI/XbaI) | This study |
| pTrc99a-*ubiB* (K407A) | pTrc99a carrying *ubiB_Ec_* (K407) (EcoRI/XbaI) | This study |
| pTrc99a-*ubiB_Fn_* (D307N) | pTrc99a carrying *ubiB_Fn_* (D307N) (EcoRI/BamHI) | This study |
| pTrc99a-*ubiB_Xc_* (D313N) | pTrc99a carrying *ubiB_Xc_* (D313N) (EcoRI/BamHI) | This study |
| pETM40-MBP-*ubiB*^ΔC47^ | pETM40 vector carrying *ubiB* gene coding for *E. coli* UbiB deleted for 47 amino acids at the C-terminal end and fused with the His_8_-maltose binding protein (Kan^R^) | This study |
| pETM40-MBP-*ubiB*^ΔC47, D310N^ | pETM40-MBP-*ubiB*^ΔC47^ encoding the D310N substitution | This study |
| pETM40-MBP-*ubiB*^ΔC47, D288N^ | pETM40-MBP-*ubiB*^ΔC47^ encoding the D288N substitution | This study |
| pETM40-MBP-*ubiB*^ΔC47, H290L^ | pETM40-MBP-*ubiB*^ΔC47^ encoding the H290L substitution | This study |
| pETM40-MBP-*ubiB*^ΔC47, N293L^ | pETM40-MBP-*ubiB*^ΔC47^ encoding the N293L substitution | This study |
| pTrc99a-MBP-*ubiB*^ΔC47^ | pTrc99a carrying MBP-*ubiB* ^Δ47^ (NcoI/XhoI) | This study |

**Table S4. Primers used in this study.**

All primers are listed from 5’ to 3’ extremities. Restriction sites are underlined and the site directed mutations are in red bold. *Xc*: *X. campestris*, *Fn*: *F. novicida*.

| **Primers** | **Sequence (5’→3’)** | **Purpose** |
| --- | --- | --- |
|  |  | **Cloning to the plasmid** |
| 5’-*ubiB_Xc_* | CCTAAGAATTCATGAAAGCCATTCTGCGC | *ubiB_Xc_* FW EcoRI |
| 3’-*ubiB_Xc_* | TTAGGGGATCCTTAACGACGCAGCCACG | *ubiB_Xc_* RV BamHI |
| 5’-*ubiB_Fn_* | TTTTGAATTCATGATTAAAAAATTTCTAAGGCTTATCTAC | *ubiB_Fn_* FW EcoRI |
| 3’-*ubiB_Fn_* | TTTTGGATCCTTATTTCTCCTTTTTAAAACTATAGTAAATC | *ubiB_Fn_* RV BamHI |
| 5’-*ubiB_Ec_* | TTTTGAATTCATGACGCCAGGTGAAGTACGG | *ubiB_Ec_* FW EcoRI |
| 3’-*ubiB_Ec_* | TTTTTCTAGATCAGCGTGTTTTGCGCCAACCG | *ubiB_Ec_* RV XbaI |
| 5’-*ubiBΔ^C47^_Ec_* | cgagaatctttattttcagggcgccatgacgccaggtgaagtac | *ubiB_Ec_* FW |
| 3’-*ubiBΔ^C47^_Ec_* | gtacctcattagtcctcgagTTACGATTGTCCCTGACG | *ubiB_Ec_* RV |
| **Primers** | **Sequence (5’→3’)** | **Site directed mutagenesis** |
| 5’-*ubiB_Xc_*-D313N | GTTTTATTGCACTG**AAT**TTTGGTATTATG | *ubiB_Xc_* (D313N) FW |
| 3’-*ubiB_Xc_*-D313N | CATAATACCAAA**ATT**CAGTGCAATAAAAC | *ubiB_Xc_* (D313N) RV |
| 5’-*ubiB_Fn_*-D307N | CCTAAATATATTTCAATA**AAC**TTTGGTATCGTTGG | *ubiB_Fn_* (D307N) FW |
| 3’-*ubiB_Fn_*-D307N | CCAACGATACCAAA**GTT**TATTGAAATATATTTAGG | *ubiB_Fn_* (D307N) RV |
| 5’-*ubiB_Ec_*-D310N | CCGAAATATATCGGCATT**AAT**TGCGGGATTGTTG | *ubiB_Ec_* (D310N) FW |
| 3’-*ubiB_Ec_*-D310N | CAACAATCCCGCA**ATT**AATGCCGATATATTTCGG | *ubiB_Ec_* (D310N) RV |
| 5’-*ubiB_Ec_*-D288N | CTTTTTCCATGCC**AAT**ATGCACCCTGGC | *ubiB_Ec_* (D288N) FW |
| 3’-*ubiB_Ec_*-D288N | GCCAGGGTGCAT**ATT**GGCATGGAAAAAG | *ubiB_Ec_* (D288N) RV |
| 5’-*ubiB_Ec_*-H286L | GACAGCTTTTTC**CTT**GCCGATATGCAC | *ubiB_Ec_* (H286L) FW |
| 3’-*ubiB_Ec_*-H286L | GTGCATATCGGC**AAG**GAAAAAGCTGTC | *ubiB_Ec_* (H286L) RV |
| 5’-*ubiB_Ec_*-H290L | CATGCCGATATG**CTC**CCTGGCAACATC | *ubiB_Ec_* (H290L) FW |
| 3’-*ubiB_Ec_*-H290L | GATGTTGCCAGG**GAG**CATATCGGCATG | *ubiB_Ec_* (H290L) RV |
| 5’-*ubiB_Ec_*-K153A | GAGGTGGTGATT**GCA**GTCATCCGCCCG | *ubiB_Ec_* (K153A) FW |
| 3’-*ubiB_Ec_*-K153A | CGGGCGGATGAC**TGC**AATCACCACCTC | *ubiB_Ec_* (K153A) RV |
| 5’-*ubiB_Ec_*-N293L | ATGCACCCTGGC**CTC**ATCTTCGTAAGC | *ubiB_Ec_* (N293L) FW |
| 3’-*ubiB_Ec_*-N293L | GCTTACGAAGAT**GAG**GCCAGGGTGCAT | *ubiB_Ec_* (N293L) RV |

**> WP_011035479.1 – 1674 bp**

gaattcATGAAAGCCATTCTGCGCGCAAGCCGTATTGGTCGTGTGATTCTGCGTTATCGTCTGGATGCACTGCTGGAAGGCACCCCGGCAGAACGTTGGCTGCGTCTGGCCAAACCGTTTGTGCCGCGTGCAAGCGCTGAAATTGCAGCCCAGAGTCGTGGTGCCCGTCTGCGCCTGGCCCTGCAGGAACTGGGTCCGATTTTTGTTAAATTTGGTCAGATTCTGTCCACCCGTCGTGATCTGATTCCGCCGGATGTGGCAGAAGAACTGACCCTGCTGCAGGATCGTGTTAAACCGTTTGATGGTGAAGCAGCCCGTCTGATTGTTGAACGTGCACTGGGTCTGCCGGTAAGCGTTGCTTTTGCTGCATTTGATACCACCCCGCTGGCATCTGCCTCAATTGCTCAGGTGCATGCCGCAACGCTGCCTCCTGATGCTAACGGTCTGCGTCGTGAAGTTGTCGTCAAAGTTCTGCGTCCGGATATTGAACGTCAGATTGATGCTGATATTGCACTGCTGCATAGCCTGGCAACCCTGGTTGAACGTACCCATCCGCGCGCAGATAAAATTCGTCCGCGTGAAGTTGTTGCCGAAATTGAAGGCACACTGAGCGCAGAACTGGATCTGCAGCGTGAAGGTGCAAATGCAAGCGTCCTGCGTCGTTTTTGGGAAGGTAGTGATGATCTGTATGTACCGGAAGTTATTTGGAGTCATACGGCAGAACGTGCACTGACACTGGAACGTGTTTATGGTATTCCGAGCGATGATGTTGCAAAACTGGATGCAGCAGGTATTGATCGTAAAGCACTGGCCGCAAAAGGCGTGCGTGTTTTTTATACCCAGGTTTTTCGCGATAATTTTTTTCATGCGGATGCACATGCGGGTAATATTTGGGTTGATAGTGATCCGGAACGTCGTCTGAATCCTCGTTTTATTGCACTGGATTTTGGTATTATGGGTCAGCTGTCACAGGAAGATCAGTATTATCTGGCAGAAAATTTTATGGCAATCTTTCATAAAGATTATCGCCGTATGGCCGAACTGCATGTTGAAGCAGGTTGGATGCCGAGCAACGTTCGTATCGATGAACTGGAAGCAGCAGCACGTAGTGTTTGTGAACCGTATTTTACCCGTCCGCTGTCCGAAATTAGCCTGGCGGAAGTCCTGATTAAACTGTTTCGTGTTGCACAGCGTTATGAACTGACCCTGCAGCCGCAGCTGATTCTGCTGCAGAAAACCCTGCTGAATATTGAAGGTGTTGGTCGTCAGCTGGACCCTAAACTGGATATCTGGGCAGTTGCTCGTCCAGTTCTGGAACGTATTCTGCGTGAACGTTATAGCCCTCGTCGTGTGCTGGGTGAACTGCGCAAACGTCTGCCGGAAATTATGACCCATGCACCTGATATGCCGCGTCTGGTTCATTCATGGCTGAAACAGCAGGTTGAAGGTCGTCATCAGCTGGATATTCGTAGCAGTGAACTGCGTGCCCTGGATCTGAGCCTGCGTAAACTGCAGACCCGTGTTGTTACCGCAATTACCGGCAGTGGTCTGCTGGTTGTTGCAGCAGTTCTGTATGGTCTGCATCCTGATGGTTGGTATCTGGGTACCGTTCCTGTTTGGAGCTGGATTTCAGGCGGTGCTGGTAGCGCAGCACTGCTGATTGCGTGGCTGCGTCGTTAAggatcc

**Dataset S2.** **Optimized sequence of *X. campestris* *ubiB* gene.** The start and stop codon are highlighted respectively in yellow and green. The added restriction enzyme sites are underlined (EcoRI at 5’end and BamHI at 3’end).

1.	Pelosi L, Vo CD, Abby SS, Loiseau L, Rascalou B, Hajj Chehade M, Faivre B, Gousse M, Chenal C, Touati N, Binet L, Cornu D, Fyfe CD, Fontecave M, Barras F, Lombard M, Pierrel F. 2019. Ubiquinone biosynthesis over the entire O_2_ range: characterization of a conserved O_2_-independent pathway. *mBio* 10.

2.	Kazemzadeh K, Hajj Chehade M, Hourdoir G, Brunet CD, Caspar Y, Loiseau L, Barras F, Pierrel F, Pelosi L. 2021. The biosynthetic pathway of ubiquinone contributes to pathogenicity of *Francisella novicida*. *J Bacteriol* 203:e0040021.

**References**
